# Asymmetric voltage attenuation in dendrites can enable hierarchical heterosynaptic plasticity

**DOI:** 10.1101/2022.07.07.499166

**Authors:** Toviah Moldwin, Menachem Kalmenson, Idan Segev

## Abstract

Long-term synaptic plasticity has been shown to be mediated via cytosolic calcium concentrations ([Ca^2+^]). Using a synaptic model which implements calcium-based long-term plasticity via two sources of Ca^2+^, NMDA receptors and voltage-gated calcium channels (VGCCs), we show in dendritic cable simulations, that the interplay between these two calcium sources can result in a diverse array of heterosynaptic effects. When spatially clustered synaptic input produces a local NMDA spike, the resulting dendritic depolarization can activate VGCCs at non-activated spines, resulting in heterosynaptic plasticity. NMDA spike activation at a given dendritic location will tend to depolarize dendritic regions that are located distally to the input site more than dendritic sites that are proximal to it. This asymmetry produces a hierarchical effect in branching dendrites, where an NMDA spike at a proximal branch can induce heterosynaptic plasticity primarily at branches that are distal to it. We also explored how simultaneously activated synaptic clusters located at different dendritic locations synergistically affect the plasticity at these locations, as well as the heterosynaptic plasticity of an inactive synapse “sandwiched” between them. We conclude that the inherent electrical asymmetry of dendritic trees enables sophisticated schemes for spatially targeted supervision of heterosynaptic plasticity.

## Introduction

The brain is believed to learn and store information via modifying the strengths of the synapses between neurons, a process known as long-term plasticity (Bliss & Collingridge, 1993; Bliss & Lomo, 1973; Hebb, 1949; Humeau & Choquet, 2019; Nabavi et al., 2014; Whitlock et al., 2006). Experimentally, plasticity can be induced via a variety of stimulation protocols (Artola et al., 1990; Bi & Poo, 1998; Bliss & Collingridge, 1993; Bliss & Lomo, 1973; O’Connor et al., 2005b; G. M. Rose & Dunwiddie, 1986; Shouval et al., 2010). While some plasticity-inducing protocols such as spike-timing-dependent plasticity (STDP) require postsynaptic depolarization (Bi & Poo, 1998), in many cases it is possible to produce long-term potentiation (LTP) or long-term depression (LTD) via presynaptic stimulation alone (e.g. using high or low frequency stimulation, respectively) (Artola et al., 1990; O’Connor et al., 2005a). Some have argued that presynaptic inputs (without post synaptic spiking activity) are the primary driver of plasticity in the hippocampus (Hardie & Spruston, 2009; J. Lisman & Spruston, 2005; White et al., 1988) as well as in some cases in the cortex (Kumar et al., 2021).

Over the past decades, since first proposed by John Lisman (J. Lisman, 1989), evidence has mounted for a calcium-based theory of plasticity, known as the calcium control hypothesis (Cho et al., 2001; Cummings et al., 1996; J. Lisman, 1989; Mulkey & Malenka, 1992; Shouval et al., 2002; Yang et al., 1999). In this framework, synapses change their strength depending on the cytosolic calcium concentration ([Ca^2+^]) at the postsynaptic dendritic spine. If the [Ca^2+^] is low, no change occurs. If the [Ca^2+^] rises above a critical threshold for depression (*θ_D_*), long-term depression (LTD) occurs and the synaptic strength is decreased. If the [Ca^2+^] is above the critical threshold for potentiation (*θ_P_*), long-term potentiation (LTP) occurs and the synaptic strength is increased (Figure 1A). (It is usually assumed that *θ_P_*>*θ_D_* for cortical and hippocampal neurons (Artola et al., 1990; J. Lisman, 1989), but the reverse may be true for cerebellar Purkinje cells (Coesmans et al., 2004; Piochon et al., 2016)). It is believed that calcium promotes LTP via pathways involving protein kinases such as calmodulin kinase (CaMKII) (J. Lisman, 1989; Malenka et al., 1989; Malinow et al., 1989; Neveu & Zucker, 1996), whereas promoting LTD via phosphatases such as calcineurin (J. Lisman, 1989; Mulkey et al., 1993, 1994).

**Figure 1.**
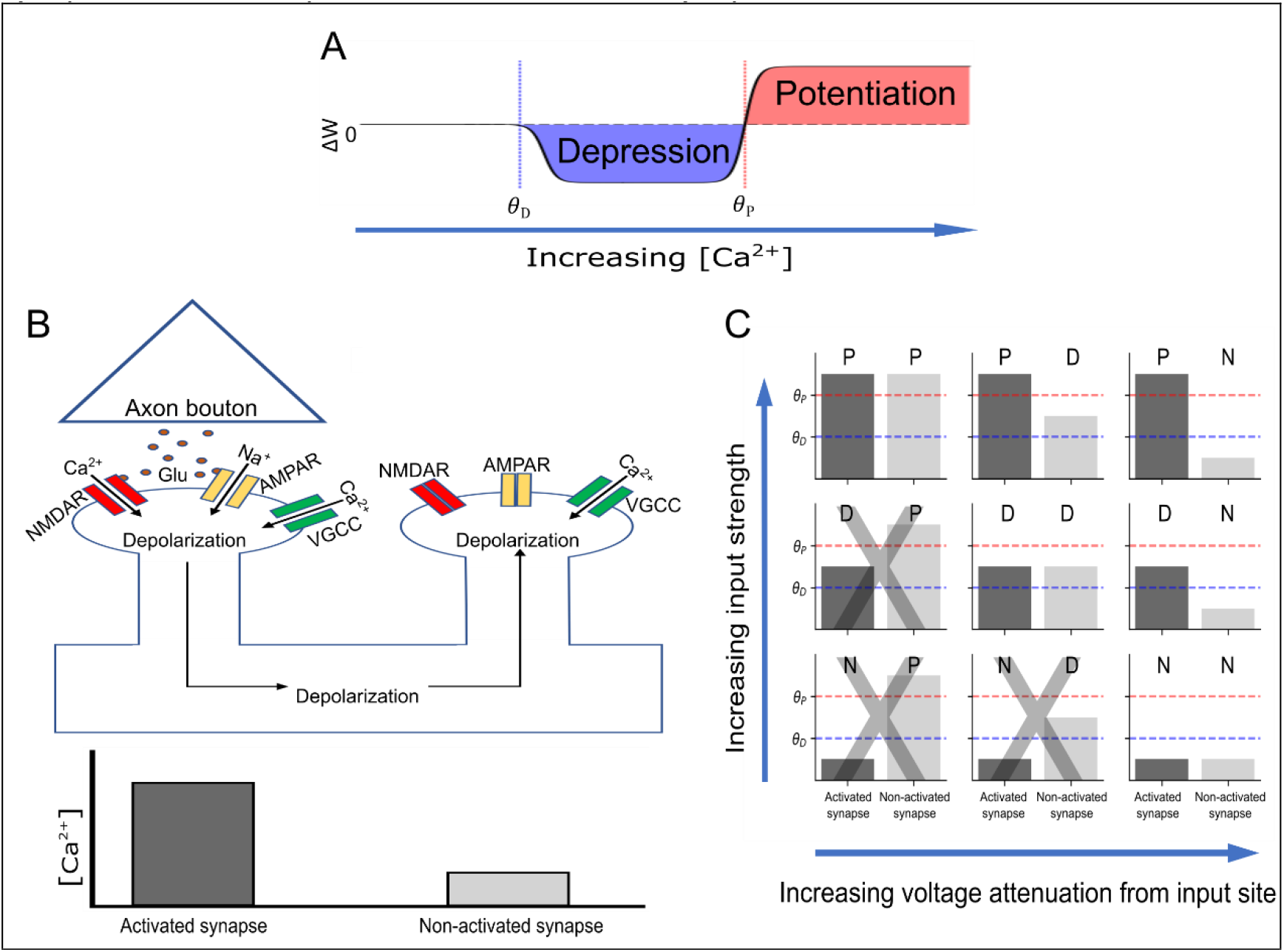
Induction of homosynaptic and heterosynaptic plasticity with NMDA receptors and VGCCs. (**A1**) Calcium control hypothesis. The synapse is weakened when the spine [Ca^2+^] crosses the depression threshold *θ_D_* and strengthened when the [Ca^2+^] crosses the potentiation threshold *θ_P_*. (**B**) VGCC hypothesis for heterosynaptic plasticity. A presynaptic neuron spikes, releasing glutamate from its axonal bouton which binds to the AMPA and NMDA receptors of the (homosynaptic) postsynaptic spine, causing calcium influx through the NMDA channel and depolarization of the spine. The depolarization opens the VGCC in the activated spine, causing additional calcium influx. The depolarization spreads and depolarizes other (heterosynaptic) spines that had not been activated, opening VGCCs in these spines. However, the overall [Ca^2+^] in the non-activated spines is smaller, as NMDA receptors were not activated. (**C**) Schematic diagram of how input strength (e.g. cluster size) and spatial voltage attenuation can affect homosynaptic and heterosynaptic plasticity. N,P, and D indicate No change, Potentiation, or Depression. Panels with a gray ‘X’ indicate scenarios that violate the assumption that activated spines have at least as much [Ca^2+^] than non-activated spines.

There are several sources of plasticity-inducing calcium at synapses. Two of the most prominent sources are the ligand- and voltage-gated NMDA receptor and the voltage gated calcium channel (VGCC). Experimentally-induced plasticity is disrupted or prevented when NMDA receptors or VGCCs are blocked, indicating that the calcium current through these sources is essential for long term plasticity (Bi & Poo, 1998; Dudek & Bear, 1992; Fino et al., 2010; Golding et al., 2002; Shindou et al., 2011). We note that internal calcium stores of calcium can also contribute to long term plasticity (Nishiyama et al., 2000; Rose and Konnerth, 2001; Royer and Paré, 2003; Jo et al., 2008; Camire and Topolnik, 2014; Evans and Blackwell, 2015; O’Hare et al., 2022), see **Discussion**.

One of the original motivations for the calcium-based plasticity theory (J. Lisman, 1989; J. E. Lisman, 2001) was the phenomenon of heterosynaptic plasticity: sometimes, when a target synapse is subjected to a plasticity protocol, other non-activated synapses are affected as well (see (Chater & Goda, 2021; Chistiakova et al., 2014) for reviews). For example, when LTP is induced at a target synapse, other synapses in the neuron can be depressed (Lynch et al., 1977). A calcium-based model can explain this phenomenon if the potentiating protocol produced a large [Ca^2+^] influx (above *θ_P_*) in the target spine, and a smaller [Ca^2+^] (above *θ_D_* but below *θ_P_*) in non-target synapses. Lisman (J. E. Lisman, 2001) proposed that this might happen in the following manner: when a target synapse is activated, such as by stimulating its presynaptic axons, NMDA receptors in the target spine are activated by the presynaptic glutamate, producing a calcium influx sufficient to potentiate the synapse. In addition to increasing the [Ca^2+^] in the target spine locally, the excitatory current also depolarizes the dendrite. If the depolarization is sufficient to activate VGCCs in other spines, they will also experience an influx of calcium, but smaller than that of the target spine, where calcium can accumulate from both from NMDA receptors and VGCCs. If the [Ca^2+^] produced by the VGCCs is above *θ_D_* but below *θ_P_*, the non-target synapses (where NMDA receptors are not activated) will depress (Figure 1C).

Heterosynaptic plasticity has also been shown to be spatially sensitive, with different plastic effects being observed at non-target synapses depending on where they are located relative to the target synapse. Some studies show heterosynaptic plasticity within short distances (~10 μm) from the target synapse (Chater & Goda, 2021; Royer & Paré, 2003; Tong et al., 2021), whereas other studies show heterosynaptic effects at up to 70 μm away from the activated synapses (Engert & Bonhoeffer, 1997) or even effects that spread from the basal to apical tree in hippocampal pyramidal neurons (Lynch et al., 1977). While the short-range effects can be potentially be explained by molecular diffusion (Chater & Goda, 2021), it is unclear what the underlying principles are that determine the spatial spread of heterosynaptic plasticity over long distances, or what the functional significance of such heterosynaptic changes might be.

Another issue that arises under the calcium control hypothesis pertains to how simultaneous synaptic input at different regions of the dendrite affects plasticity. It is known that NMDA synapses can interact synergistically such that when multiple nearby synapses are activated simultaneously, the observed somatic EPSP is larger than the linear sum of individual EPSPs, due to the voltage-dependence of the NMDA receptor (Polsky et al., 2004). However, it was not systematically explored how simultaneous synaptic activity at different locations on the dendrite affect plastic changes at both activated and non-activated synapses.

Recently, a model synapse was developed as part of the Blue Brain Project (Chindemi, Abdellah, Amsalem, Benavides-Piccione, Delattre, Doron, Ecker, King, Kumbhar, Monney, Perin, Rössert, Geit, et al., 2020) which incorporates NMDA receptors, VGCCs, and calcium-dependent long-term plasticity dynamics. This synapse model (with some modifications described below in **Methods**) enables us to explore Lisman’s hypothesis about the calcium basis of heterosynaptic plasticity in a dendritic cable model, which provides insight into the spatial properties of heterosynaptic plasticity.

## Results

### Possible Heterosynaptic Effects

We begin by considering the range of possible heterosynaptic effects that may occur according to the hypothesis that homosynaptic plastic effects from presynaptic plasticity-induction protocols are induced by calcium influx from both NMDA receptors and VGCCs, whereas heterosynaptic effects are induced only via calcium influx through VGCCs. In this view, a spine activated with presynaptic input will almost inevitably have a higher calcium concentration than non-activated spines, as both NMDA receptors and VGCCs can enable calcium influx in the activated spine, but only VGCCs can be opened in the non-activated spine. (Figure 1B).

Because calcium thresholds for LTP and LTD can vary from cell to cell (Yang et al., 1999), and can also be changed via meta-plastic processes (Abraham & Bear, 1996), we generically map out several possible results that can occur to an activated and non-activated synapse given a few basic assumptions: 1) an activated synapse has a higher calcium influx than a non-activated synapse; 2) plasticity thresholds and voltage-gated channel densities are approximately the same from spine to spine within the same neuron and 3) that the [Ca^2+^] threshold for potentiation is higher than that of depression, i.e. *θ_P_* > *θ_D_*. We also disregard the magnitude of the plastic change and only consider the direction (potentiation or depression), as we assume that after inducing plasticity, synapses eventually drift toward a binary potentiated or depressed state, based on (Graupner & Brunel, 2012).

Given these assumptions, the following plastic effects can result (Figure 1B): If the activated synapse is potentiated, non-activated synapses can also be potentiated (PP), but they can also be depressed (PD), or undergo no change (PN). If the activated synapse is depressed, non-activated synapses can also be depressed (DD) or undergo no change (DN). If the activated synapse does not change, neither will the non-activated synapse (NN). Given assumptions our assumptions above, the following possibilities are *not* possible: DP, NP, and ND (Figure 1C). (In the event that *θ_D_* > *θ_P_*, as in Purkinje cells, the allowed possibilities are PP, PN, DP, DD, DN, NN, and the disallowed possibilities are PD, NP, and ND, however the simulations used in this study assume *θ_P_* > *θ_D_* to match known results from the hippocampus and cortex (Cho et al., 2001; Mulkey & Malenka, 1992). We also assume that the input to the activated synapse is not sufficiently large to put it into a post-potentiative neutral zone where the calcium concentration is so high that potentiation mechanisms are inactivated, see (Tigaret et al., 2016)).

Intuitively, given values for *θ_D_* and *θ_P_*, homosynaptic effects from presynaptic input protocols will vary based on the “strength” of synaptic input (e.g. input frequency or cluster size, as we use here). Strong inputs can potentiate the synapse, medium strength inputs can depress it, and weak inputs will induce no change. As heterosynaptic effects are mediated by dendritic depolarization, heterosynaptic effects will also depend on the input strength to the activated synapse, in addition to other factors which determine the “spillover effect” of the activated synapse’s dendritic depolarization on non-active synapses.

One important factor that can affect this electrical spillover between synapses is the distance of the non-activated synapse from the activated synapse. Because voltage attenuates with electrotonic distance in dendrites (Rall, 1967; Rall & Rinzel, 1973), we would naively expect that non-activated synapses which are closer to the activated synapse will see more dendritic depolarization, and are thus likely to have a larger calcium influx through VGCCs, than synapses that are further away from the activated synapse.

### Asymmetric attenuation of EPSPs and NMDA dendritic spikes in dendrites

The distance-dependent attenuation description of heterosynaptic plasticity is complicated by the fact that voltage attenuation in the dendrite is highly asymmetric. For distal dendritic inputs, the proximal dendrites and soma act as a current “sink”. This gives rise to a strong asymmetry in voltage attenuation in dendrites (Rall & Rinzel, 1973).

To demonstrate the effects of dendritic location on voltage attenuation, we created a ball-and-stick cable model with a 200 μm long cylindrical cable coupled to an isopotential soma (Figure 2A). We enlarged the diameter of the soma to replicate the electrical sink effect that would occur in a layer 5 cortical pyramidal neuron with a full dendritic morphology (Hay et al., 2011) (see Methods, Supplementary 1). We placed a cluster of 5 dendritic spines with excitatory synapses at 100 μm from the soma. We also placed non-activated spines at 60 μm and 200 μm from the soma. We simultaneously activated all synapses in the spine cluster located at 100 μm from the soma and recorded the local voltage at the soma, the dendrite and at the heads of both the activated and non-activated spines. Voltage attenuated slightly from the heads of the activated spines to the spine base, but almost no attenuation was visible from the base of the activated spines at 100 μm to the head of the distal non-active spine at 200 μm. By contrast, the voltage attenuated substantially from the base of the activated spines to the base of the proximal non-active spine (Fig. 2B, see also (Segev & Rall, 1988)). Qualitatively, substantial attenuation from spine to dendrite but not from dendrite to spine is consistent with recent experimental work(Cornejo et al., 2021).

**Figure 2.**
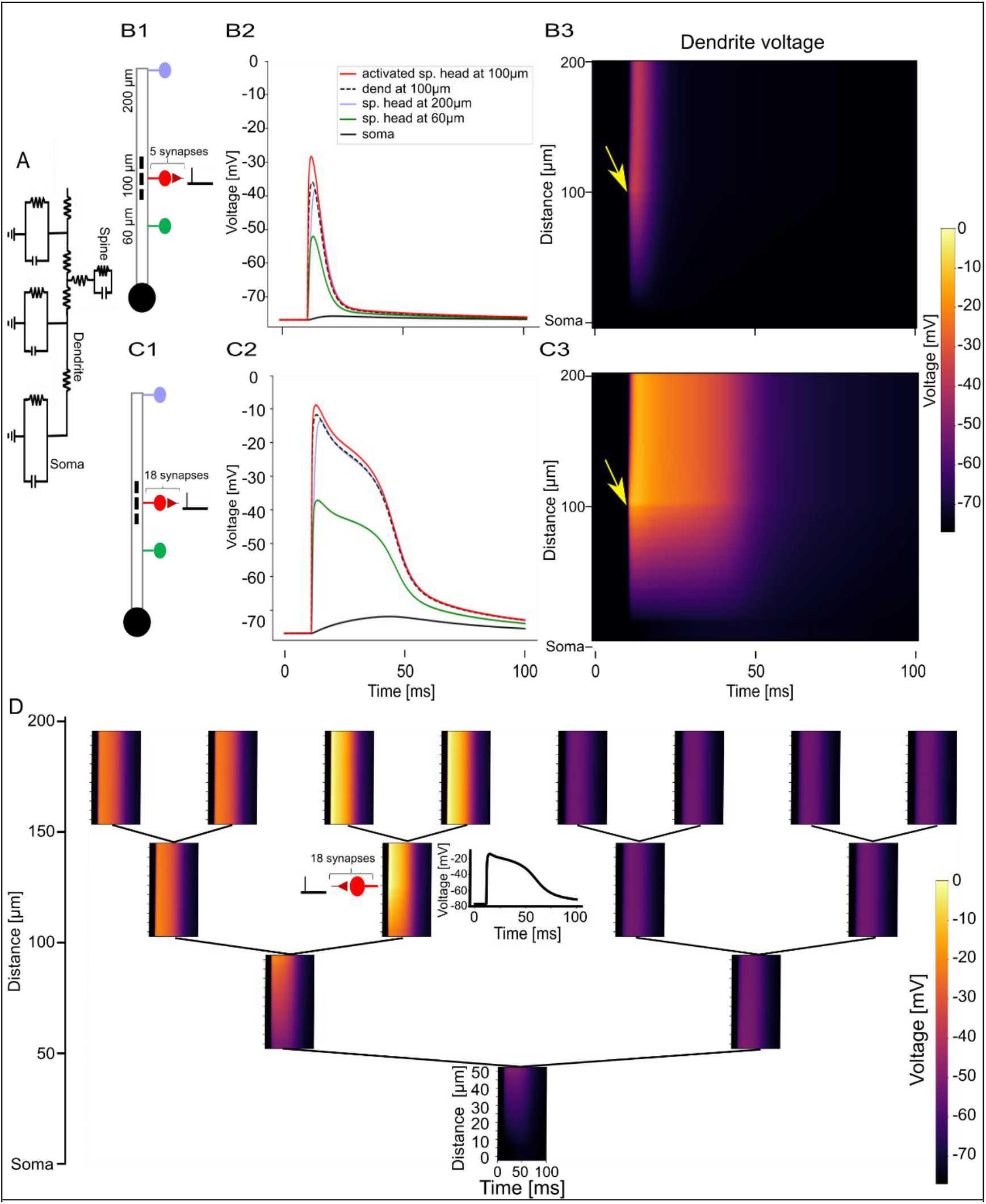
Asymmetric attenuation of EPSPs and NMDA spikes in dendritic cable models. (**A**) Circuit diagram of a ball and stick dendritic model with spines. (**B1**) Experiment schematic. A cluster of 5 spine synapses located at 100 μm from the soma are simultaneously activated. Voltage is recorded from one of the activated spine heads (red spine), its base (black dashed line), a non-activated spine at 200 μm from the soma (blue spine) and a non-activated spine at 60 μm from the soma (green spine) and the soma (black solid circle). (**B2**) Voltage traces from recording sites depicted in B1. Voltage is largest at the activated spine head (red solid line); it attenuates somewhat to the spine base (dashed black line). Very little attenuation occurs from the activated spine base to the distal spine head (blue), but significant attenuation is observed toward the proximal spines (green line) and the soma (black line). (**B3**) Voltage recordings along the dendrite during the experiment depicted in B1. Color depicts dendritic voltage as a function of time (horizontal axis) and distance from soma (vertical axis). Arrow indicates time and location of the activated synaptic cluster. (**C1-C3**) Same as B1-B3 except a cluster of 18 synapses are simultaneously activated to generate an NMDA spike. (**D**) Dendritic voltage heatmaps in each branch of an order-3 branching dendritic model in response to an NMDA spike initiated via activating a cluster of 18 synapses at the indicated dendritic location. Inset: voltage trace at the base of the activated spine cluster.

We replicated this experiment with a cluster of 18 synapses at 100 μm from the soma, which was sufficient to generate an NMDA spike in these spines (see also (Eyal, Verhoog, Testa-Silva, Deitcher, Benavides-Piccione, et al., 2018)). (We note that it is possible to create an NMDA spike with fewer clustered synapses if the synapses are activated at a high frequency (Dembrow & Spain, 2022; Polsky et al., 2009), however in this work, for simplicity, we only vary the cluster size.) The same asymmetric effect as described above was qualitatively observed for the NMDA spike; voltage attenuation was very minor from the activation site to the distal tip and very substantial from the activation site toward the soma (Figure 2B-C).

We next demonstrated how the asymmetric attenuation manifests in a branching dendrite model. We created an order-3 branching dendritic tree (i.e. 3 bifurcating branching levels emanating from the 0^th^-level main branch) coupled to a soma compartment (Figure 2D). We simultaneously activated a cluster of 18 synapses in the middle of a branch at the 2^nd^ level branch, generating a local NMDA spike there. The NMDA spike propagated to the distal daughter branches with minimal attenuation, propagated to the sister branches of the activated branch and their daughter branches with mild attenuation, and propagated to the rest of the dendritic tree with substantial attenuation (Figure 2D). The stark contrast in the depolarization magnitude of different regions of the dendritic tree in response to a local NMDA spike raises the possibility that asymmetric voltage attenuation may play a functional role in governing plasticity processes in different parts of the dendritic tree.

### Asymmetric voltage attenuation produces asymmetric heterosynaptic plasticity

To explore how asymmetric voltage attenuation can impact heterosynaptic plasticity we placed a spine at each segment of the ball and stick model shown in Fig. 3A1 (one spine every 10 μm) and activated a cluster of 27 synapses at the center of the dendrite. This produced a large NMDA spike at the activated spines, depolarizing the dendrite sufficiently to open VGCCs at non-activated spines (Fig. 3A4).

**Figure 3.**
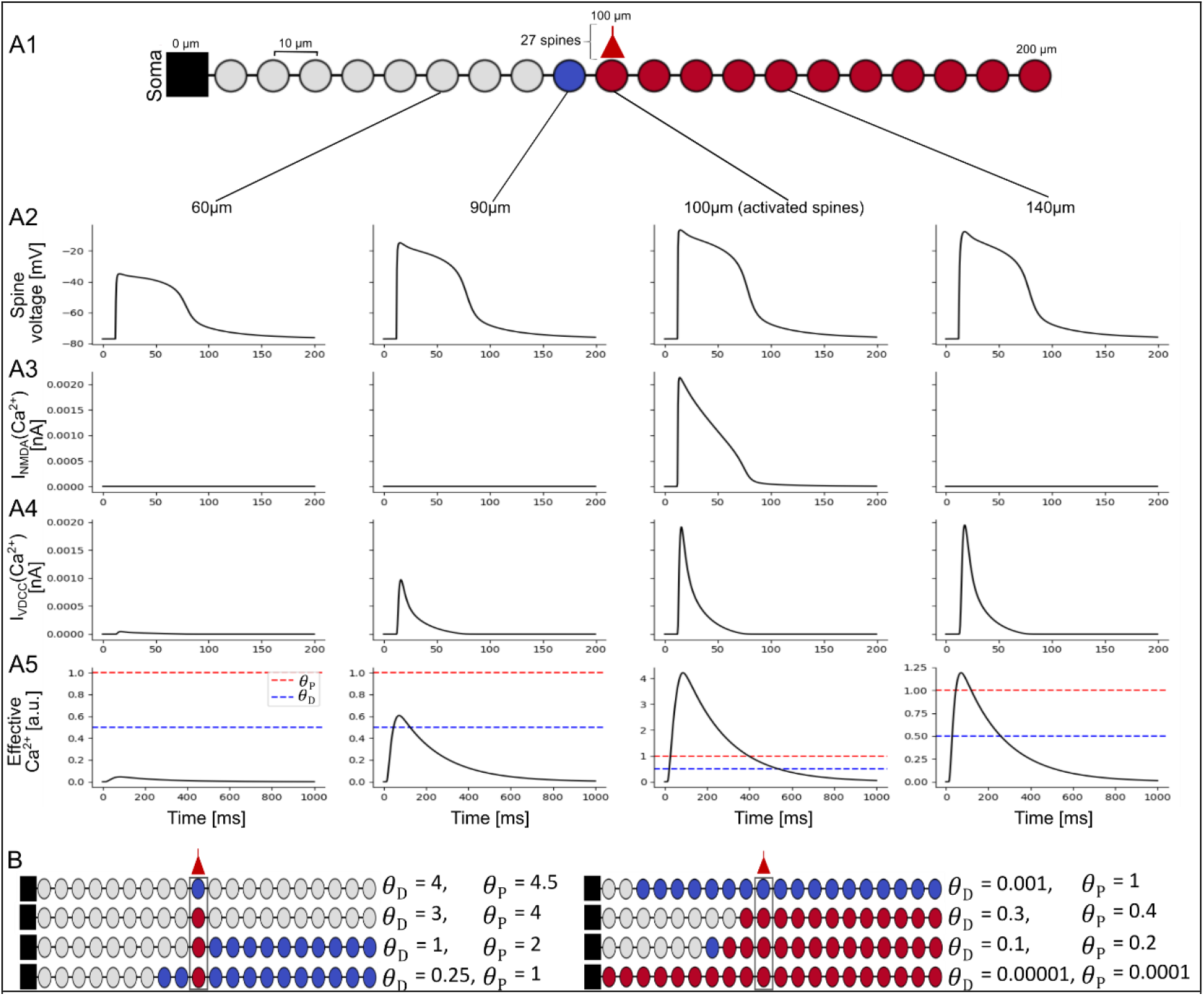
Asymmetric heterosynaptic plasticity induced by VGCCs. **(A1)** Top: A ball and stick model dendrite with spines (circles) placed every 10 μm. A cluster of 27 spines (shown as a single circle with an input) is activated at the center of the dendrite (100 μm), generating an NMDA spike which results in homosynaptic and heterosynaptic plasticity. The activated spines and the spines distal to it are potentiated (red), the spine 10 μm proximal to the activated spine is depressed (blue), and the other proximal spines do not change (gray). (**A2**) Spine head voltage traces shown at 60 μm, 90 μm, 100 μm (exemplar activated spine), and 140 μm. The NMDA spike is seen at all spines, but the voltage at the proximal location (60 μm) is substantially attenuated. (**A3**) Ca^2+^ current through the NMDA receptor at the 4 depicted spines. Only the activated spine has NMDA current because NMDA receptors are ligand gated. (**A4**) Ca^2+^ current through the VGCCs at the depicted spines; Ca^2+^ current depends on local voltage (from A2). (**A5**) Effective [Ca^2+^] (as accumulated by the Ca^2+^ integrator) at the depicted spines. At 60 μm the [Ca^2+^] is below *θ_D_* (blue dashed line) so no change occurs, at 90 μm, the [Ca^2+^] reaches above *θ_D_* but below *θ_P_* (red dashed line) so depression occurs, at 100 μm and at 140 μm the [Ca^2+^] reaches above *θ_P_*, so the synapses are potentiated. (**B**) As in A1 (27 synapses activated at 100 μm) but with different calcium thresholds, resulting in different heterosynaptic effects.

At the active site, both NMDA channels and VGCCs provided a substantial amount of calcium current into the cell, allowing [Ca^2+^] to surpass *θ_P_*, generating homosynaptic potentiation. The depolarization spreading from the activated spines was sufficient to open VGCCs at distal spines, providing enough calcium current to induce heterosynaptic potentiation. However, at a spine located 10 μm proximal to the input site, the voltage had already attenuated sufficiently such that the [Ca^2+^] there following the opening of the local VGCCs induced depression (Fig. 3A1, 3A5). At 20 μm proximal to the input site, the voltage attenuated such that the [Ca^2+^] from the VGCCs was insufficient to cross *θ_D_*, so that the synapses on this spine and all other proximal spines were left unchanged (Figure 3A).

Clearly, the specific plasticity outcomes that we observed hold true only for the specific calcium thresholds for plasticity used in our simulation. We therefore performed the same experiment with a variety of different values for the plasticity thresholds. We observe that it is generally easier to induce heterosynaptic plasticity at spines that are distal to the activation site rather than proximal to it (Figure 3B).

This directional asymmetry in heterosynaptic effects is especially pronounced when considering a branching dendrite, as asymmetric attenuation from an input site can create branch-dependent dendritic depolarization (Rall & Rinzel, 1973). To demonstrate this effect, we placed spines every 10 μm on the order-3 branched model described above. We activated clusters of 20, 30, 40, or 50 synapses at the 0^th^-3^rd^-order branches and observed the plastic changes at all modeled spines (Figure 4).

**Figure 4.**
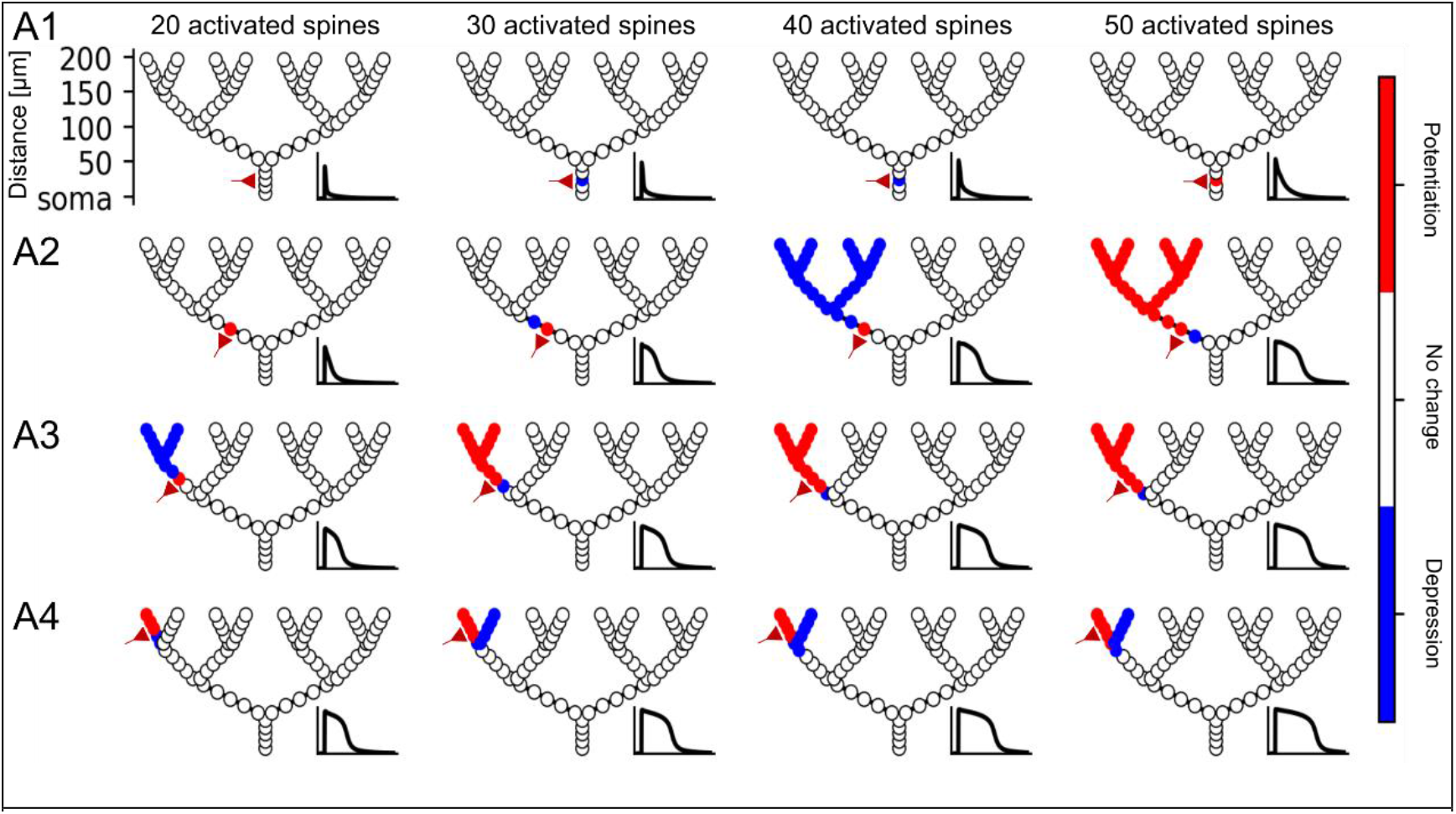
Hierarchical heterosynaptic plasticity in a branching dendritic model. **(A1)** A synaptic cluster of 20 (first column), 30 (second column), 40 (third column), or 50 (fourth column) spines are simultaneously activated at the indicated location (schematic red synapse) on the proximal 0^th^-order branch. Insets: spine head voltage at the activated sites. Due to the low input resistance at the proximal branch, 20 synapses are insufficient to generate homosynaptic plasticity, while 30 and 40 synapses generate local homosynaptic depression (blue), and 50 synapses generate homosynaptic potentiation (red). **(A2)** Same as (A1), but the activated cluster is now at the 1^st^-order branch. 20 activated synapses create homosynaptic potentiation, 30 or 40 activated synapses create a short-duration NMDA spike and heterosynaptic depression at synapses distal to the activation site, whereas 50 activated synapses create a prolonged NMDA spike and heterosynaptic potentiation at distal sites. **(A3)** Activated cluster at the 2^nd^-order branch; 20 activated synapses cause a short NMDA spike and distal heterosynaptic depression, 30 or more synapses create a prolonged NMDA spike and heterosynaptic potentiation at distal synapses. **(A4)** Activated cluster at 3^rd^-order branch. 20 activated synapses are sufficient to cause distal heterosynaptic potentiation and 30 or more activated synapses also produce heterosynaptic depression at the sister branch to the input site.

If the activated synaptic cluster is placed on the 0^th^-order branch emerging from the soma, due to the relatively low input resistance there, it is difficult to induce an NMDA spike or homosynaptic plasticity (30 spines are required for depression, 50 for potentiation). Even 50 synapses are insufficient in this model to produce an NMDA spike at the most proximal branch, and thus heterosynaptic plasticity is not induced (Fig. 4A1). If we move the cluster up to a 1^st^-order branch, homosynaptic potentiation is induced with 20 synapses, and an NMDA spike is elicited with 30 synapses. When 40 spines are activated, spines on the dendritic tree distal to the input site are depressed, and when 50 spines are activated, these distal synapses are potentiated (Fig. 4A2). When placed at a 2^nd^-order branch, 20 activated synapses are already sufficient to produce an NMDA spike and heterosynaptic depression at dendritic spines distal to the input site, and 30 spines turns the heterosynaptic depression to potentiation (Fig. 4A3). At the 3^rd^-order branch, the input resistance is sufficiently large to give rise to both homosynaptic and heterosynaptic potentiation at spines distal to the input site with 20 synapses, and with 30 synapses the voltage propagates sufficiently in the proximal direction to depress the input site’s sister branches.

The tiered nature of heterosynaptic plasticity in dendrites, where proximal inputs can induce heterosynaptic plasticity at branches that are distal to it, suggests that dendritic branches might supervise each other in a hierarchical manner. Branches that are closer to the soma (although not so close that it is difficult to generate an NMDA spike) can “teach” the branches that are distal to it because the NMDA spike preferentially propagates backwards toward distal locations, leading to heterosynaptic potentiation or depression in descendant branches. Moreover, distal branches with high input resistances may be able to supervise plasticity in their sibling branches via a competitive process where an input sufficient to depress the branch with homosynaptic input will heterosynaptically depress synapses on a sibling branch.

### Synergistic Synaptic “Sandwiching”

Until now, we have only looked at heterosynaptic effects produced by the activation of a single cluster of co-localized spines, generating a single local NMDA spike. It is possible that multiple clusters can be activated simultaneously, generating diverse depolarization effects in the dendritic tree (Palmer et al., 2014). From a plasticity standpoint, it is important to think about how the clusters of activated synapses can affect each other (through both VGCC and NMDA-dependent activations) as well as how they affect inactive synapses via heterosynaptic plasticity (through VGCC activation). While it is not feasible to explore the full combinatorial space of cluster activations, we consider a canonical case in our ball-and-stick model where an inactive spine, placed 130 μm from the soma, is “sandwiched” in between two spine clusters, one proximal (60 μm from the soma) and one distal (200 μm from the soma, Fig. 5A). This case is important for understanding the mechanisms governing heterosynaptic plasticity because it illustrates the tradeoff between two principles. On the one hand, voltage attenuates more steeply toward the soma. On the other hand, it is easier to generate a large/prolonged NMDA spike at distal synapses, due to the higher input resistance at distal locations (Doron et al., 2017; Poleg-Polsky, 2015).

**Figure 5.**
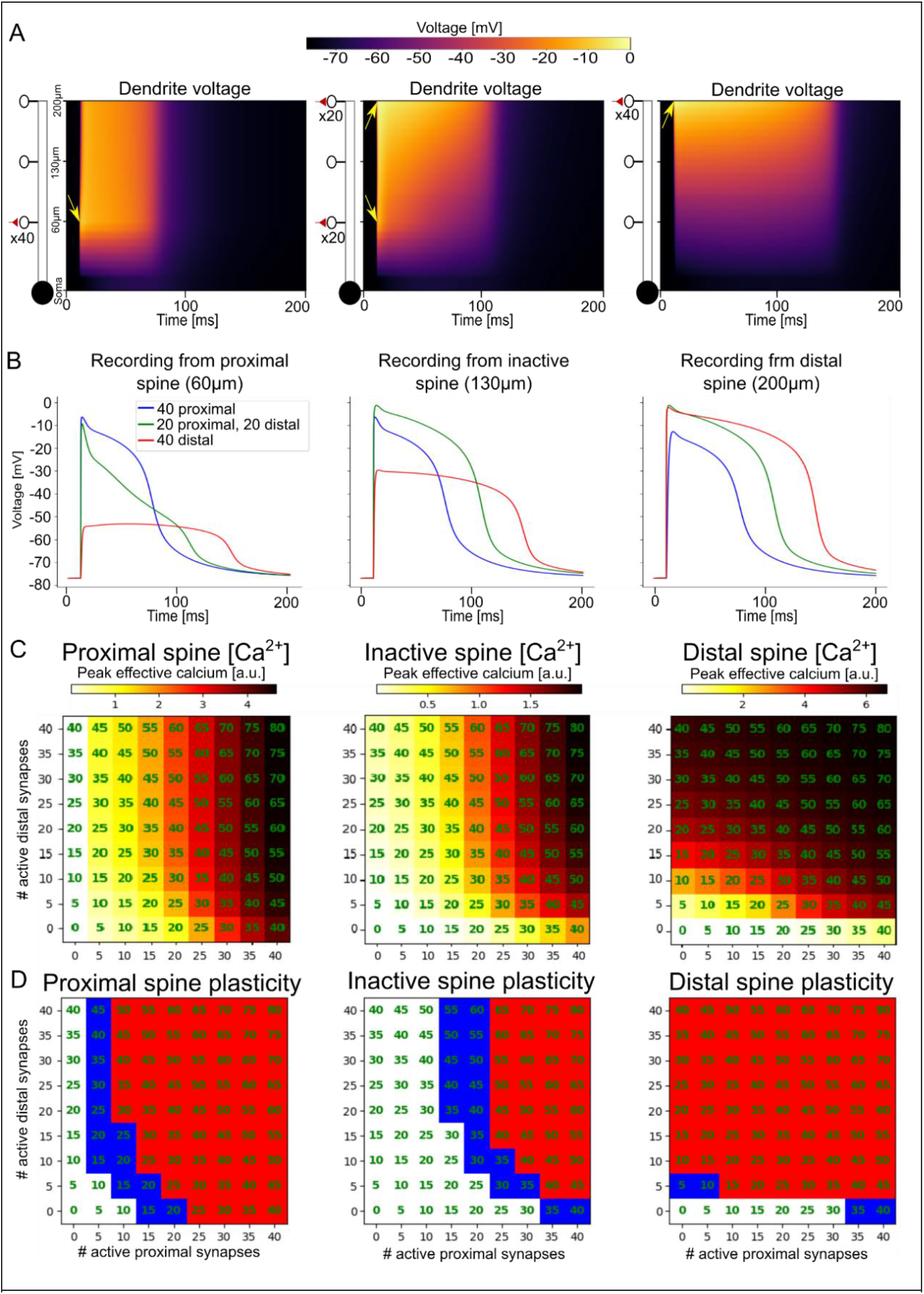
Proximal and distal clusters can create synergistic heterosynaptic effects for “sandwiched” synapses. **(A)** Dendritic voltage over time in a ball-and-stick model (inset at left) when 40 synapses are placed proximally (left) distally (right) or distributed evenly between the proximal and distal locations (center). Arrows show the location and time of the activated synapses. **(B)** Spine head voltage recordings from a proximal activated spine (left), the central non-activated spine (center), and a distal activated spine (right) for the 3 cases shown in A. **(C)** Peak calcium influx at the proximal (left), central (middle), or distal spine (right) as a heatmap, for different numbers of spines placed in the distal and proximal clusters. Annotations indicate total number of spines (proximal + distal). **(D)** Plastic effect (red: potentiation, blue: depression, white: no change) due to the calcium influx from (C).

To illustrate this tradeoff, suppose we have 40 active synapses to distribute between the proximal cluster and the distal cluster with the goal of maximizing the depolarization, and thus the heterosynaptic calcium influx, at the centrally located inactive synapse. If the input resistance effect dominates, it would be better to place all synapses distally. If the asymmetric voltage attenuation effect dominates, we might assign all 40 synapses to the proximal cluster. In fact, however, it seems that the answer lies in between these two extremes: Placing 20 synapses each at the proximal and distal location result in a slightly larger depolarization at the heterosynaptic synapse than placing all 40 synapses together in a single cluster at either the proximal or distal location (Figure 5A-B). The synergy between distal and proximal clusters is not restricted to the case of 40 synapses; for any given number of synapses there appears to be a “sweet spot” for distributing those synapses between the proximal and distal locations to maximize heterosynaptic effects at the central location, albeit with a tendency to assign more synapses to the proximal location (Figure 5C-D).

The increased depolarization when the synapses are divided in separate clusters can be explained by the fact that there are diminishing returns for placing additional synapses at the same location, due to the reduced driving force when the dendrite is depolarized to near its reversal potential. It is thus better to separate the synapses into separate clusters at locations that are somewhat electrically separated to avoid “wasting” synapses on a dendritic segment that is already maximally depolarized. (Additional synapses can still increase the *duration* of an NMDA spike when the branch is depolarized to near its electrical reversal, however for the purposes of heterosynaptic plasticity it is often crucial to maximize the peak voltage at the heterosynaptic synapse in to ensure that the peak calcium through the VGCCs passes the plasticity thresholds. The duration above the plasticity thresholds also affects the magnitude of the plastic changes in the early phase of long-term plasticity, but in the bistable Graupner-Brunel bistable model, magnitude information is lost after several hours in the late phase of plasticity when the synaptic weights are stabilized into a binary UP/DOWN state, see (Graupner & Brunel, 2012).)

The benefit of dividing synapses into two groups is not observed at the proximal and distal locations themselves. While active proximal synapses do increase calcium influx at distal synapses and vice versa (due to both NMDA and VGCC voltage-dependence), to maximize peak calcium influx at proximal spines, it is best to put all the synapses proximally, and to maximize peak calcium influx at distal spines, it’s best to put all the synapses distally (Figure 5C-D).

The synergistic heterosynaptic sandwiching effect also pertains in a branched neuron model. We placed varying numbers of activated spines on a proximal branch (second layer) and a distal branch (fourth layer) in our 4-layer branched model to observe the heterosynaptic effects at a non-activated spine on the central branch (third layer). As in the ball-and stick model, the peak calcium at the non-active, central spine was maximized when active synapses were distributed between the proximal and distal branch (Figure 6A-C, Supplementary Figure S2).

**Figure 6.**
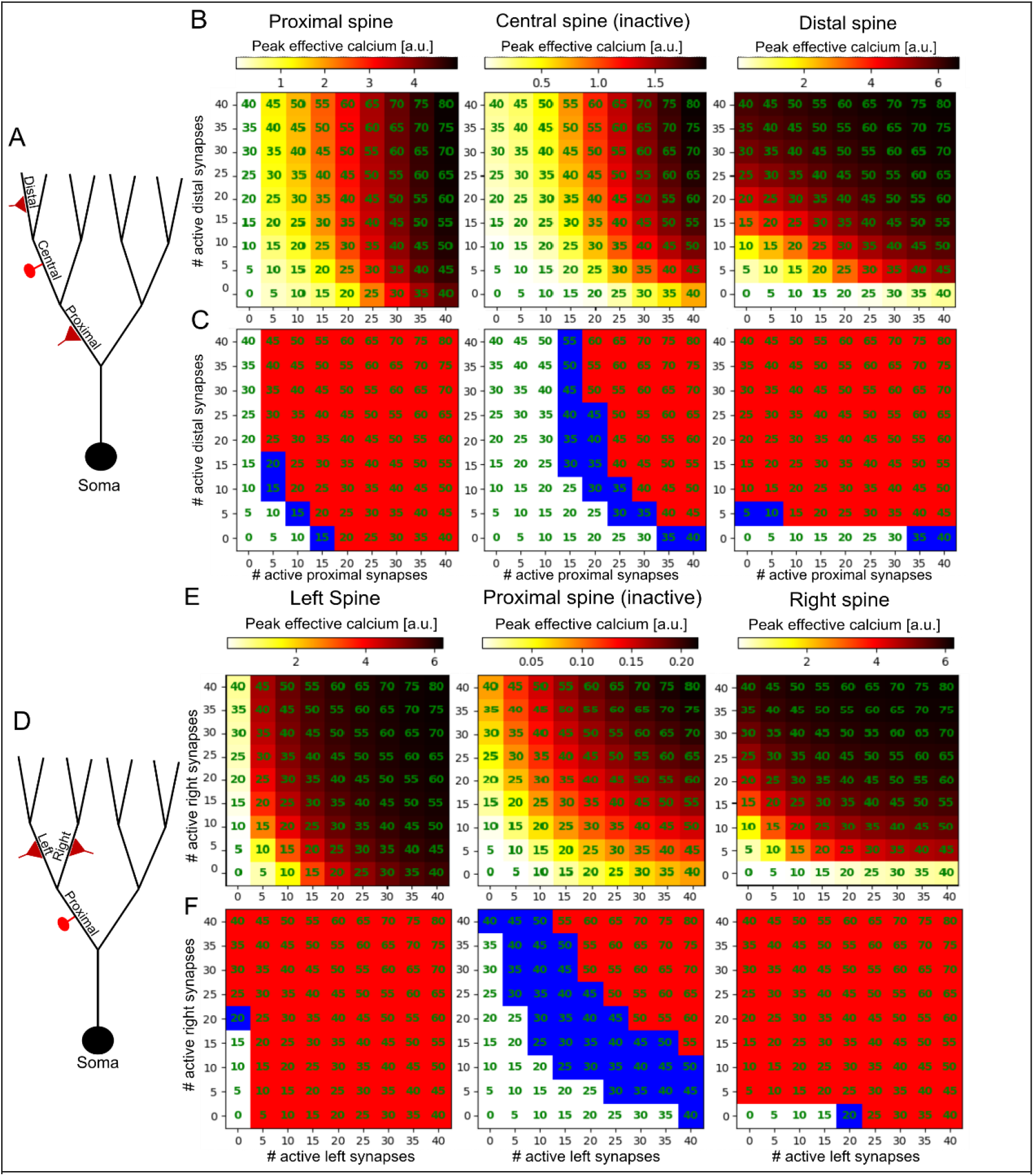
Vertical and horizontal heterosynaptic “sandwiching” in a branched dendritic model. **(A)** Experiment schematic for “vertical sandwiching” in a branched model. Clusters of spines are activated at proximal (1^st^ order) and distal (3^rd^ order) branches to explore their effect of on each other and on a non-activated spine located at a “central” branch (2^nd^ order) in between them. **(B)** Peak calcium concentration (A.U) at an exemplar spine on the proximal (left), central (middle) or distal (right) branches. **(C)** Plastic effect on each spine as a function of cluster sizes (red: potentiation, blue: depression, white: no change). **(D)** Experiment schematic for “horizontal sandwiching” in a branched model. Clusters of spines are activated at 2^nd^-order left and right branches to explore how clusters on sister branches affect each other as well as a non-activated synapse on their parent (1^st^ order) branch. **(E-F)** Calcium and plasticity for the horizontal sandwiching experiment as in **C-D.** See Supplementary Figures S2 and S3 for voltage traces and heatmaps.

In additional to this “vertical sandwiching” scenario, we also explore a “horizontal sandwiching” case, where an inactive spine is placed in the middle of a branch at the second branching layer, and varying numbers of active synapses are placed at its left and right daughter branches at the third branching layer. We again observe in this context that from the perspective of the inactive spine on the proximal parent branch, dividing the active spines between the left and right daughter branches tends to maximize the peak calcium available for producing heterosynaptic plasticity. As we would expect from the symmetry of the left and right branches relative to the parent branch, the peak heterosynaptic calcium tends to be maximal when the left and right branches have the same number of activated spines (Figure 6D-F, Supplementary Figure S3).

We have thus shown that when an inactive synapse is placed between two synaptic clusters, whether it is “vertically sandwiched” between a distal and proximal branch or “horizontally sandwiched” between two of its daughter branches, plasticity-inducing calcium influx tends to be greater than if all the active synapses were placed in a single cluster. This raises the possibility that in addition to the hierarchical supervision effect we showed above, it may be possible to engineer synapse placement in a sophisticated manner to maximize heterosynaptic plasticity induction without requiring an excessive number of active synapses at the same location.

## Discussion

Our simulations have shown a wide range of consequences for synaptic plasticity arising from the hypothesis that heterosynaptic plasticity might result from dendritic depolarization-induced calcium influx through VGCCs. Simple dendritic cable models, combined with model synapses containing NMDA and VGCCs channels, were sufficient to produce spatially-sensitive heterosynaptic plasticity effects using a standard calcium-based plasticity mechanism. Specifically, we have demonstrated that a strong dendritic input which generates an NMDA spike can induce heterosynaptic plasticity at dendritic sites that are distal to the input due to asymmetric voltage attenuation in dendrites. This asymmetry can create a hierarchical heterosynaptic effect in a branching dendrite structure, whereby clustered inputs to a branch closer to the soma can act as “supervisors” to synapses located on more distal branches. Moreover, when two input clusters are active, each cluster can increase the plasticity-inducing calcium influx at spines in the other cluster, as well as at non-active spines. Additionally, calcium influx to a non-active spine can be maximized by dividing activated spines into two clusters, rather than placing all activated spines at the same location.

The extent to which these phenomena occur in biology remains an open question, and we encourage experimentalists to use the predictions of our model to design experiments to test whether hierarchical heterosynaptic plastic effects indeed occur in the brain. If our predictions are borne out by experiments, then heterosynaptic plasticity can produce a richer repertoire of plastic effects than have been previously considered. If dendritic NMDA spikes indeed act as heterosynaptic supervisors for other (more distal) synapses, the dendritic branching structure and the location of NMDA spike induction become essential for plasticity induction. Location-sensitive NMDA spike-dependent plasticity rules are particularly critical in light of findings that backpropagating somatic action potentials may not reach distal synapses, and therefore do not induce plasticity in these synapses, whereas NMDA spikes can induce plasticity there (Kumar et al., 2021), see also (Hardie & Spruston, 2009; J. Lisman & Spruston, 2005).

Further work can explore diverse neuronal types with different dendritic morphologies to examine whether the branching structure of different neurons may lend themselves to different kinds of plasticity computations. For example, the elaborate fractal branching structure of Purkinje neurons may lend those neurons to be optimized for segregated hierarchical units (see (Liu et al., 2016; Piochon et al., 2007, 2010) regarding the presence of NMDARs in Purkinje neurons and their influence on plasticity). Conversely, neurons with sufficiently long, branching dendrites (such as apical dendrites of L2/3 cells) may exhibit more attenuation in the proximal-to-distal direction and thus behave less hierarchically, as sufficiently long distal branches themselves can act as electrical sinks relative to the parent branch (Landau et al., 2022).

The branch-dependent variation of heterosynaptic plasticity we show in our model is in line with the theory that the dendritic branch may be a fundamental computational unit in the neuron (Branco & Häusser, 2010; Koch et al., 1983; Segev & Rall, 1998). Consistent with the idea that neurons can behave as a two-layer and even multiple layer neural network (Beniaguev et al., 2021; Poirazi et al., 2003; Poirazi & Mel, 2001), hierarchical plasticity can potentially serve as a biophysical basis for a multi-layer learning algorithm within a single neuron, perhaps akin to the backpropagation algorithm in deep neural networks in feed-forward artificial neural networks (Jones & Kording, 2021; Rumelhart et al., 1986). The details of how such an algorithm would operate remain an open avenue for investigation.

The hierarchical plasticity phenomenon as suggested here is complicated somewhat by our “sandwiching” results, which demonstrate that, for a given number of activated synapses, heterosynaptic effects can be maximized by distributing them into two (or possibly more) spatially segregated clusters instead of placing them all at the same location. This points to the possibility of an even more sophisticated supervision scheme, where multiple synaptic clusters can be strategically placed at different dendritic locations to produce spatially targeted heterosynaptic plasticity. Spatiotemporally-targeted inhibition may also help shape the spread of heterosynaptic plasticity. Further experimental and theoretical work could explore these possibilities in more detail. In any event, the diverse heterosynaptic effects we have shown here provide support for the claim that neurons may behave as complex nonlinear units (Beniaguev et al., 2021; Jones & Kording, 2021; Koch & Segev, 2000; Larkum, 2022; Poirazi et al., 2003) as opposed to simple perceptrons where synapses are modified independently (Moldwin & Segev, 2020). Moreover, the pronounced asymmetrical voltage attenuation in dendrites and the attendant consequences for heterosynaptic plasticity shown in our simulations indicate that computational models which make use of distance-dependent NMDA superlinearities (e.g. (Mel, 1991; Moldwin et al., 2021)) should take into account branching structure and synaptic location relative to the soma in addition to the relative distance of synapses from each other.

### Additional biological considerations

Our model, in line with the proposal of Lisman (J. E. Lisman, 2001), assumes that the only medium of communication between active and inactive synapses is dendritic voltage depolarization, which can activate VGCCs of other nonactivated dendritic spines. We note that many other mechanisms for the induction of heterosynaptic plasticity have been suggested (See (Chater & Goda, 2021; Chistiakova et al., 2014) for reviews). One alternative possibility is that calcium itself diffuses from one synapse to another, however experimental evidence suggests that calcium diffusion from the spine head into the dendritic shaft is negligible (Sabatini et al., 2002; Yuste & Denk, 1995). Other molecules have also been implicated in inducing heterosynaptic effects, such as h-Ras, Rac1, RhoA, Arc, BDNF-TrkB, CaMKII and calcineurin (Chater & Goda, 2021; Tong et al., 2021), however these molecules have only been shown to diffuse up to 10μm along the dendrite, while heterosynaptic effects have been shown to occur at much larger distances between activated and non-activated synapses (Engert & Bonhoeffer, 1997; Lynch et al., 1977). As such, the depolarization-based model remains an important candidate mechanism of heterosynaptic plasticity. It may be that there are different short-distance and long-distance heterosynaptic effects, with short-distance effects occurring via molecular mechanisms such as local CamKII and Calcneurin activity, while long-distance effects may be due to the voltage mechanism we describe here.

Regarding the fidelity of the parameters our simulation to biological reality, there are several questions that would require additional experimental evidence and more detailed models to fully confirm. Our calcium channel model assumed a single type of calcium channel, and we chose a conductance value that roughly corresponds to what might expect as the aggregate conductance of all high-voltage activated VGCCs. A more precise model that includes all forms of VGCCs with their appropriate unitary conductances, kinetics, and densities would allow to examine our claims with greater precision. Additionally, the kinetics of calcium accumulation and the plasticity thresholds for calcium used here could be better constrained with more experimental evidence. We also assumed that calcium channel density and plasticity thresholds were the same from spine to spine; in biology these may differ on a spine-by-spine basis even with a single neuron. Moreover, the spatial effects we observed in our simulations assume a passive dendritic cable; active mechanisms in biological dendrites such as voltage-gated sodium, calcium and potassium channels have been shown to differentially modulate voltage propagation in different neurons (Golding et al., 2001), so these mechanisms would consequently be expected to modify the spatial dynamics of heterosynaptic plasticity as well.

Inhibitory synapses also likely play an important role in the spatial reach of heterosynaptic plasticity. Inhibition can have different consequences for the neuronal voltage depending on the location of the inhibitory synapses (Bar-Ilan et al., 2012; Gidon & Segev, 2012; M. Jadi et al., 2012; M. P. Jadi et al., 2014) as well as their timing relative to excitatory NMDA inputs (Doron et al., 2017). As such, spatiotemporally targeted inhibition can be used to modulate the heterosynaptic effects we describe here, enabling a bidirectional control system for heterosynaptic plasticity.

Several studies have shown that internal calcium stores play an important role in both homosynaptic and heterosynaptic plasticity (Camire & Topolnik, 2014; Evans & Blackwell, 2015; Jo et al., 2008; Nishiyama et al., 2000; O’Hare et al., 2022; C. R. Rose & Konnerth, 2001; Royer & Paré, 2003). Ryanodine and IP3 receptors produce calcium-induced calcium release (CICR) which can affect plasticity in a variety of ways. Although experimental results regarding the role of CICR are more subtle than the model we have presented here, one way to think about CICR is as an amplifier of calcium coming from NMDA and voltage gated calcium channels. As such, assuming that CICR increases monotonically with calcium from extracellular sources, the basic qualitative principle that activated synapses will experience more calcium release than non-activated synapses due to NMDA receptor activation still holds, but it may shift the homosynaptic and heterosynaptic effects observed (e.g. from homosynaptic depression and no heterosynaptic effect to homosynaptic potentiation and heterosynaptic depression, see figure 1C). However, there is evidence that the effect of internal calcium stores is highly localized into micro domains and depends on various second messengers, resulting in a more complex picture of homosynaptic and heterosynaptic effects (Evans & Blackwell, 2015).

Another biological mechanism that may affect the plastic results we predict here are small-conductance Ca^2+^-activated K+channels (SK channels). SK channels can repolarize the membrane in response to calcium influx (Adelman et al., 2012; Rodrigues et al., 2021; Tigaret et al., 2016), potentially reducing both homosynaptic and heterosynaptic effects.

One additional crucial biological question is whether the early stage heterosynaptic plasticity induced by calcium influx is stabilized into late-term plasticity via protein synthesis, which has been shown to be necessary to make plastic changes last longer than an hour (Barco et al., 2008; Frey & Morris, 1997; Redondo et al., 2010). One recent study (Sun et al., 2021), showed that plasticity-induced protein synthesis may primarily occur within 3 μm of potentiated synapses, suggested that heterosynaptic effects may not necessarily be long-lived. However, it is possible that a very strong clustered stimulation, such as we described here, may induce protein synthesis at more distant locations.

### Methods

Simulations were done using NEURON with a Python wrapper (Carnevale & M.L. Hines, 1997; Hines et al., 2009). Code was written using Python 3.7.6 and NERUON version 7.7.2. Figures were made with the Matplotlib package.

Model parameters can be found in Table 1. The dendrite models were largely based on the layer 5 pyramidal cell model of (Hay et al., 2011) except where described otherwise in Table 1. The dendritic axial resistance and diameter were chosen such as to fit with empirically described results (Yuste & Denk, 1995)and to ensure that a robust NMDA spike could be obtained with ~20 synapses (Eyal, Verhoog, Testa-Silva, Deitcher, Benavides-Piccione, et al., 2018)The ball and stick model had a dendrite 200 μm, composed of 50 electrical segments (cylinders). The branched model had 4 levels of bifurcating branches. Each branch was 50 μm long and was composed of 10 segments.

**Table 1.**

| **Model parameters** |  |  |  |
| --- | --- | --- | --- |
|  | **Property** | **Value** | **Reference** |
| | $R_a$ | Dend: 150 $\Omega\text{cm}$ | |
| | $C_m$ | Soma: 1 $\mu\text{F}/\text{cm}^2$ | (Hay et al., 2011) |
| | | Dendrite: = 1* $\mu\text{F}/\text{cm}^2$ | (Hay et al., 2011) |
| | $E_{\text{pas}}$ | -77 mV | (Hay et al., 2011) |
| | $R_m$ | Soma: 30 $\text{K}\Omega\text{cm}^2$ | (Hay et al., 2011) |
| | | Dendrite: = 44* $\text{K}\Omega\text{cm}^2$ | (Hay et al., 2011) |
| | $g_{\text{max}}$ | AMPA (INITIAL): 1.5 nS<br>AMPA (UP): 2 nS<br>AMPA (DOWN): 1 nS | (Behabadi et al., 2012) |
|  |  | VGCC: 20 pS | (Fisher et al., 1990;<br>Snutch et al., 2013;<br>Tsien et al., 1988) |
|  |  | NMDAR: 1.31 nS | (Eyal, Verhoog, Testa-Silva, Deitcher, Piccione, et al., 2018) |
| **Morphological parameters** | Diameter | Dendrite: 0.75 $\mu\text{m}$ | (Araya et al., 2006) |
| | | Soma: 718 $\mu\text{m}$ | (Hay et al., 2011), see **Methods** and **S1** |
| | Length | Dendrite (ball and stick): 200 $\mu\text{m}$ | |
| | | (Branched model): 50 $\mu\text{m}$ per branch | |
| | | Soma: 23 $\mu\text{m}$ | (Hay et al., 2011) |
| **Spine parameters** | $R_a$ | 150 $\Omega\text{cm}$ | |
| | $R_m$ | 10.7 $\text{K}\Omega\text{cm}^2$ | (Hay et al., 2011)<br>(no spine comp.) |
| | Diameter | Head: 0.4 $\mu\text{m}$ | (Konur et al., 2003) |
| | | Neck: 0.07 $\mu\text{m}$ | Arellano et al. 2007a<br>(fit to ensure $R_{\text{neck}}$ of 226.6 $\text{M}\Omega$) |
| | Length | Neck: 0.66 $\mu\text{m}$ | Arellano et al. 2007a |
| | $R_{\text{neck}}$ | Neck: 226.6 $\text{M}\Omega$ | (Cornejo et al., 2021) |

\* $R_m$ used in the simulation was divided by 2 and $C_m$ multiplied by 2 to compensate for surface area of unmodeled spines while maintaining the membrane time constant (Eyal, Verhoog, Testa-Silva, Deitcher, Benavides-Piccione, et al., 2018)

To create a ball-and-stick model that replicates the electrical sink effect observed in the soma of a neuron with a full complement of extended dendrites, we expanded the diameter of the ball-and-stick soma such as to have the same input resistance by applying the formula 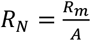, where *R_N_* is the somatic input resistance of the full layer 5 pyramidal cell model in units of Ω, *R_m_* is the membrane resistivity in units of Ωcm2, and *A* is the area of the compensated soma in units of cm2, resulting in a compensated soma diameter of 718 μm. See Supplementary 1A for a comparison of transfer resistances between the fully pyramidal model and our ball-and-stick model.

As in the Hay model, effective dendritic membrane resistivity *dendritic R_m_* is divided by two to compensate for the surface area of (unmodeled) spines and dendritic membrane capacitance *dendritic C_m_* was doubled to ensure that the membrane time constant *τ* does not change.

For synapses, we used the Blue Brain Project’s synapse model with NMDA receptors, VGCCs, and calcium-based long-term plasticity model (Chindemi, Abdellah, Amsalem, Benavides-Piccione, Delattre, Doron, Ecker, King, Kumbhar, Monney, Perin, Rössert, van Geit, et al., 2020); details of the synaptic and calcium dynamics can be found there. The calcium-based plasticity model itself is based on (Graupner & Brunel, 2012).

We made several modifications to the Blue Brain synapse: (1) We initialized the synapses in a neutral state of 1.5 nS (equivalent to *ρ* = 0.5, i.e. the unstable fixed point in the Graupner-Brunel model), so synapses could be easily depressed or potentiated when the calcium accumulator crossed the plasticity thresholds. (2) We changed the maximum conductance of the NMDA receptor (gmax_NMDA) to 1.31 nS based on (Eyal, Verhoog, Testa-Silva, Deitcher, Piccione, et al., 2018) (3) We increased the unitary conductance of the VGCCs to 20 pS based on (Fisher et al., 1990; Snutch et al., 2013; Tsien et al., 1988). While the kinetics of the VGCC model are based on the R-type VGCC, 20 pS was chosen as a rough estimate of the total unitary conductance over all high-voltage activated channels. (4) We modified the equations for total spine calcium conductance and concentration to account for the fact that we were explicitly modeling the spine as a cylinder with the parameters in Table 1.

Except where indicated, for Figures 3–6, plasticity thresholds for the [Ca^2+^] were: *θ_D_* = 0.5, *θ_P_* = 1. For Figure 7, plasticity thresholds were *θ_D_* = 0.2, *θ_P_* = 0.4. Plasticity thresholds can vary from cell to cell, so all plasticity results presented here should be taken as qualitative illustrations of possible plastic effects rather than specific quantitative predictions.

## Acknowledgements

The authors would like to thank András Ecker and Giuseppe Chindemi for the use of their plastic synapse model (Chindemi, Abdellah, Amsalem, Benavides-Piccione, Delattre, Doron, Ecker, King, Kumbhar, Monney, Perin, Rössert, van Geit, et al., 2020), their support and helpful discussions in dealing with technical and conceptual issues over the course of this project, and for their helpful comments our manuscript. We also thank Yuri Rodrigues and Thomas Chater for their thorough comments on the manuscript. This work received generous support from the Drahi family foundation, the ETH domain for the Blue Brain Project, the Gatsby Charitable Foundation, and the NIH Grant Agreement U01MH114812.

---

**Supplementary Figure S1:**
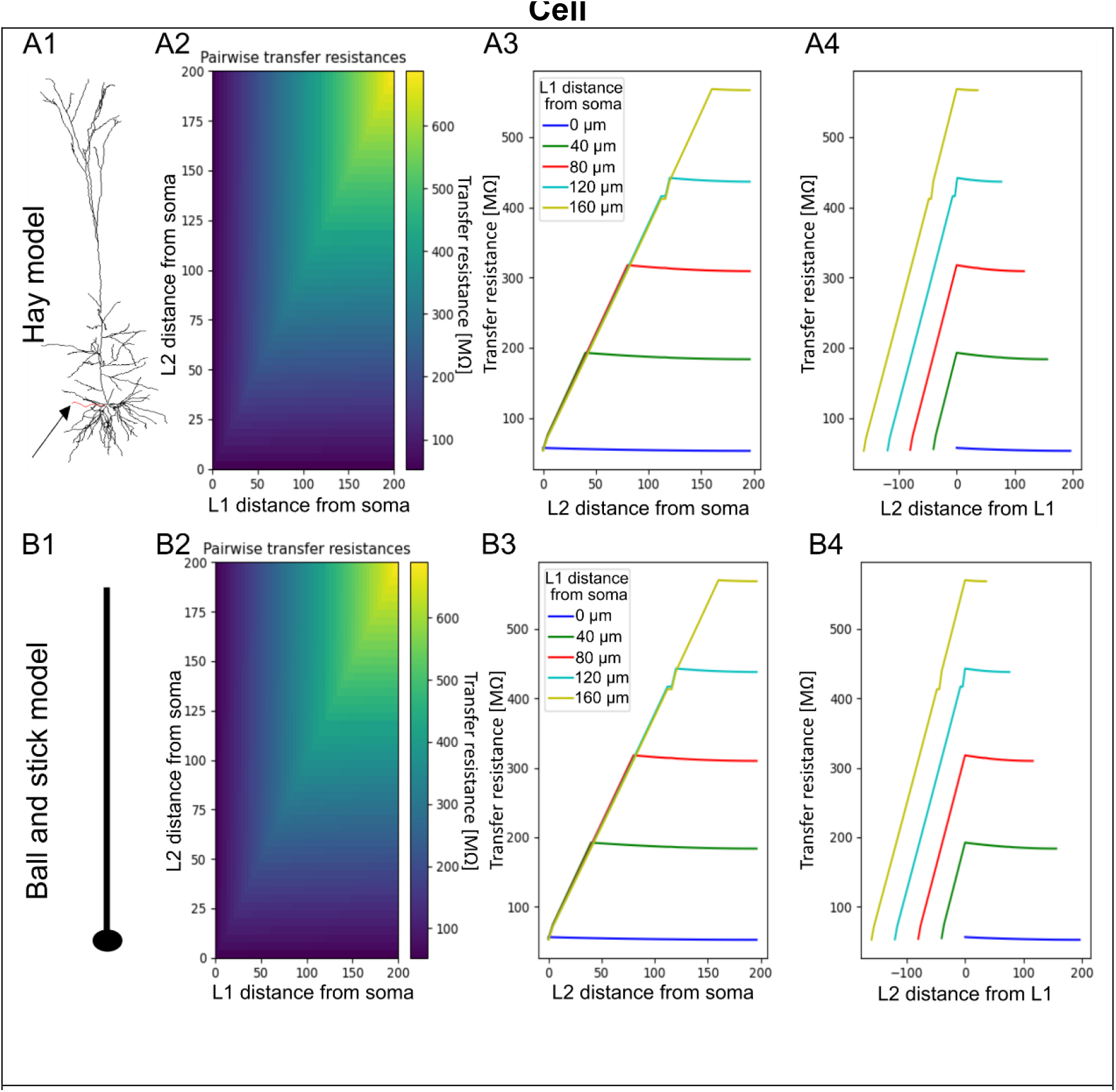
Asymmetric transfer resistances of L5PC and ball-and-stick models. (**A1**) Model of layer 5 pyramidal cell from (Hay et al., 2011). Arrow indicates a dendrite modified to have the same morphological and electrical parameters as the idealized ball-and-stick model. (**A2**) Pairwise transfer resistances between each dendritic segment, presented as heatmap. L1: location 1, L2: location 2. (**A3**) Transfer resistances from selected dendritic locations (L1) to all other dendritic locations (L2) as a function of distance of L2 from the soma. (**A4**) As is A3, except L2 is depicted as a function of distance from L1, demonstrating asymmetric attenuation from each location. Negative numbers indicate L2 is more proximal than L1, positive numbers indicate L1 (**B1-B4**) Same as A1-A4, but for the ball-and-stick model used in the paper with the soma diameter adjusted to match the somatic input resistance (*R_N_*) from the Hay L5PC model.

**Supplementary Figure S2.**
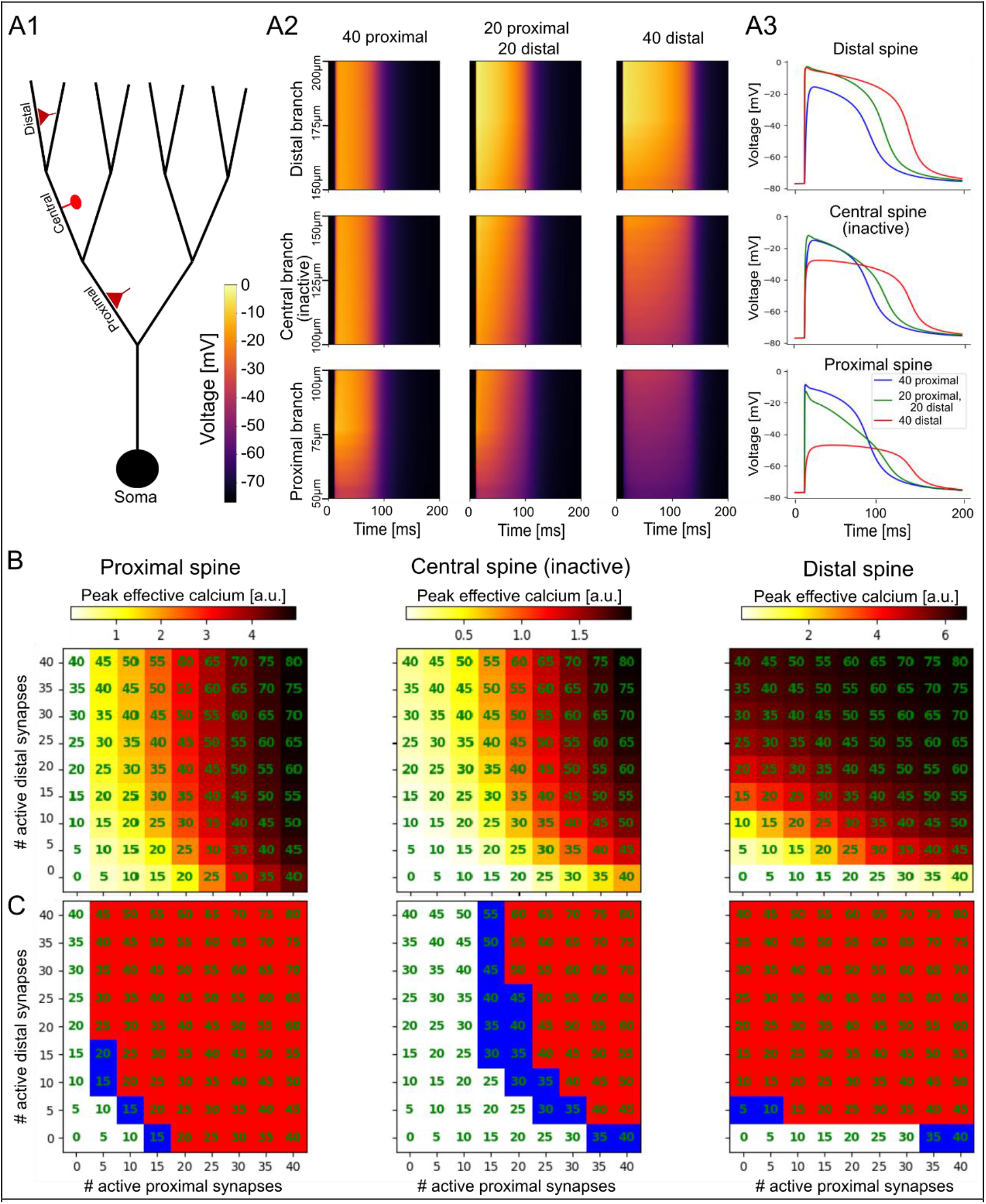
Vertical heterosynaptic sandwiching in a branched model. **(A1)** Experiment schematic. Clusters of spines are activated at proximal (2^nd^ layer) and distal (4^th^ layer) branches to explore the effect on non-activated spine on a central branch (3^rd^ layer). **(A2)** Spatiotemporal voltage profiles at the distal (top row), central (middle row), and proximal (bottom row) branches in the cases where a cluster of 40 active synapses are all placed at the center of the proximal branch (left), distal branch (right), or where two clusters of 20 synapses each are placed on the proximal and distal branches, respectively. **(A3)** Voltage traces at an exemplar spine head from the distal cluster (top), the inactive synapse on the central branch (center), or the proximal cluster (bottom) for each of the experimental protocols (40 proximal, 40 distal, 20 proximal + 20 distal). **(B)** Peak calcium at an exemplar spine on the proximal (left), central (middle) or distal (right) branches, as in Figure 5C. **(C)** Plastic effect on each spine as a function of cluster sizes (red: potentiation, blue: depression, white: no change). **B-C** same as Figure 6 in the main text.

**Supplementary Figure S3.**
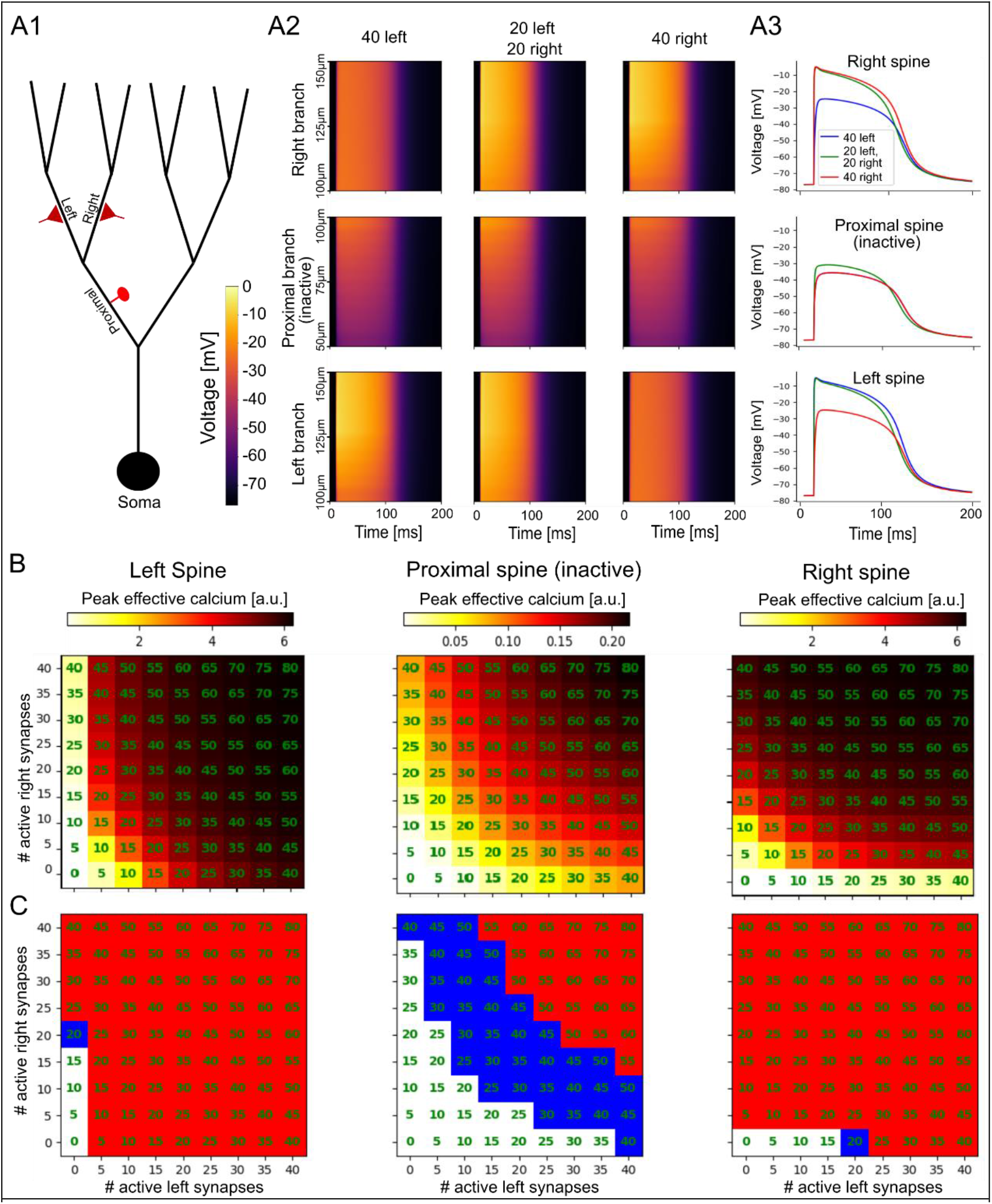
Horizontal heterosynaptic sandwiching in a branched model. **(A1)** Experiment schematic. Clusters of spines are activated at the third branching layer on the left and right branches to explore the effect on non-activated spine on a proximal parent branch (2^nd^ layer). **(A2)** Spatiotemporal voltage profiles at the right (top row), proximal (middle row), and left (bottom row) branches in the cases where a cluster of 40 active synapses are all placed at the center of the left branch (left), right branch (right), or where two clusters of 20 synapses each are placed on the left and right branches, respectively (center). **(A3)** Voltage traces at an exemplar spine head from the right cluster (top) proximal cluster (left) or the inactive synapse on the proximal parent branch (center) for each of the experimental protocols (40 proximal, 40 distal, 20 proximal + 20 distal). **(B)** Peak calcium at an exemplar spine on the left, proximal parent, or right branches, as in Figure 6B. **(C)** Plastic effect on each spine as a function of cluster sizes (red: potentiation, blue: depression, white: no change). [Note that to demonstrate plastic effects, calcium thresholds for plasticity used in this figure were different than in other figures, see **Methods**.) **B-C** same as Figure 6 in the main text.

## References

Abraham, W. C., & Bear, M. F. (1996). Metaplasticity: The plasticity of synaptic plasticity. Trends in Neurosciences, 19(4), 126–130. https://doi.org/10.1016/S0166-2236(96)80018-X

Adelman, J. P., Maylie, J., & Sah, P. (2012). Small-conductance Ca 2+-activated K + channels: Form and function. Annual Review of Physiology, 74, 245–269. https://doi.org/10.1146/ANNUREV-PHYSIOL-020911-153336

Araya, R., Eisenthal, K. B., & Yuste, R. (2006). Dendritic spines linearize the summation of excitatory potentials. www.pnas.orgcgidoi10.1073pnas.0609225103

Artola, A., Bröcher, S., & Singer, W. (1990). Different voltage-dependent thresholds for inducing long-term depression and long-term potentiation in slices of rat visual cortex. Nature, 347(6288), 69–72. https://doi.org/10.1038/347069A0

Barco, A., Lopez de Armentia, M., & Alarcon, J. M. (2008). Synapse-specific stabilization of plasticity processes: The synaptic tagging and capture hypothesis revisited 10 years later. Neuroscience & Biobehavioral Reviews, 32(4), 831–851. https://doi.org/10.1016/J.NEUBIOREV.2008.01.002

Bar-Ilan, L., Gidon, A., & Segev, I. (2012). The role of dendritic inhibition in shaping the plasticity of excitatory synapses. Frontiers in Neural Circuits, DEC. https://doi.org/10.3389/FNCIR.2012.00118

Behabadi, B. F., Polsky, A., Jadi, M., Schiller, J., & Mel, B. W. (2012). Location-dependent excitatory synaptic interactions in pyramidal neuron dendrites. PLoS Computational Biology. https://doi.org/10.1371/journal.pcbi.1002599

Beniaguev, D., Segev, I., & London, M. (2021). Single cortical neurons as deep artificial neural networks. Neuron, 109(17), 2727–2739.e3. https://doi.org/10.1016/J.NEURON.2021.07.002

Bi, G., & Poo, M. (1998). Synaptic Modifications in Cultured Hippocampal Neurons: Dependence on Spike Timing, Synaptic Strength, and Postsynaptic Cell Type. Journal of Neuroscience, 18(24), 10464–10472. https://doi.org/10.1523/JNEUROSCI.18-24-10464.1998

Bliss, T., & Collingridge, G. (1993). A synaptic model of memory: long-term potentiation in the hippocampus. Nature, 361(6407), 31–39. https://doi.org/10.1038/361031a0

Bliss, T., & Lomo, T. (1973). Long-lasting potentiation of synaptic transmission in the dentate area of the anaesthetized rabbit following stimulation of the perforant path. The Journal of Physiology, 232(2), 331–356. https://doi.org/10.1113/jphysiol.1973.sp010273

Branco, T., & Häusser, M. (2010). The single dendritic branch as a fundamental functional unit in the nervous system. Current Opinion in Neurobiology, 20(4), 494–502. https://doi.org/10.1016/j.conb.2010.07.009

Camire, O., & Topolnik, L. (2014). Dendritic Calcium Nonlinearities Switch the Direction of Synaptic Plasticity in Fast-Spiking Interneurons. Journal of Neuroscience, 34(11), 3864–3877. https://doi.org/10.1523/JNEUROSCI.2253-13.2014

Carnevale, N. T., & M.L. Hines. (1997). The NEURON Simulation Environment. Neural Computation, 1209, 1–26.

Chater, T. E., & Goda, Y. (2021). My Neighbour Hetero — deconstructing the mechanisms underlying heterosynaptic plasticity. Current Opinion in Neurobiology, 67, 106–114. https://doi.org/10.1016/J.CONB.2020.10.007

Chindemi, G., Abdellah, M., Amsalem, O., Benavides-Piccione, R., Delattre, V., Doron, M., Ecker, A., King, J., Kumbhar, P., Monney, C., Perin, R., Rössert, C., Geit, W. Van, DeFelipe, J., Graupner, M., Segev, I., Markram, H., & Muller, E. (2020). A calciumbased plasticity model predicts long-term potentiation and depression in the neocortex. BioRxiv, 2020.04.19.043117. https://doi.org/10.1101/2020.04.19.043117

Chindemi, G., Abdellah, M., Amsalem, O., Benavides-Piccione, R., Delattre, V., Doron, M., Ecker, A., King, J., Kumbhar, P., Monney, C., Perin, R., Rössert, C., van Geit, W., DeFelipe, J., Graupner, M., Segev, I., Markram, H., Muller, E., Geit, W. van,… Muller, E. (2020). A calcium-based plasticity model predicts long-term potentiation and depression in the neocortex. BioRxiv, 2020.04.19.043117. https://doi.org/10.1101/2020.04.19.043117

Chistiakova, M., Bannon, N. M., Bazhenov, M., & Volgushev, M. (2014). Heterosynaptic Plasticity. The Neuroscientist, 20(5), 483–498. https://doi.org/10.1177/1073858414529829

Cho, K., Aggleton, J. P., Brown, M. W., & Bashir, Z. I. (2001). An experimental test of the role of postsynaptic calcium levels in determining synaptic strength using perirhinal cortex of rat. The Journal of Physiology, 532(Pt 2), 459. https://doi.org/10.1111/J.1469-7793.2001.0459F.X

Coesmans, M., Weber, J. T., de Zeeuw, C. I., & Hansel, C. (2004). Bidirectional Parallel Fiber Plasticity in the Cerebellum under Climbing Fiber Control. Neuron, 44(4), 691–700. https://doi.org/10.1016/j.neuron.2004.10.031

Cornejo, V. H., Ofer, N., & Yuste, R. (2021). Voltage compartmentalization in dendritic spines in vivo. Science. https://doi.org/10.1126/SCIENCE.ABG0501

Cummings, J. A., Mulkey, R. M., Nicoll, R. A., & Malenka, R. C. (1996). Ca2+ Signaling Requirements for Long-Term Depression in the Hippocampus. Neuron, 16(4), 825–833. https://doi.org/10.1016/S0896-6273(00)80102-6

Dembrow, N. C., & Spain, W. J. (2022). Input rate encoding and gain control in dendrites of neocortical pyramidal neurons. Cell Reports, 38(7), 110382. https://doi.org/10.1016/J.CELREP.2022.110382

Doron, M., Chindemi, G., Muller, E., Markram, H., Segev, I., Eilif Muller, H. M., Segev, I., Muller, E., Markram, H., Segev, I., Eilif Muller, H. M., & Segev, I. (2017). Timed Synaptic Inhibition Shapes NMDA Spikes, Influencing Local Dendritic Processing and Global I/O Properties of Cortical Neurons. Cell Reports, 21(6), 1550–1561. https://doi.org/10.1016/j.celrep.2017.10.035

Dudek, S. M., & Bear, M. F. (1992). Homosynaptic long-term depression in area CA1 of hippocampus and effects of N-methyl-D-aspartate receptor blockade. Proceedings of the National Academy of Sciences, 89(10), 4363–4367. https://doi.org/10.1073/pnas.89.10.4363

Engert, F., & Bonhoeffer, T. (1997). Synapse specificity of long-term potentiation breaks down at short distances. Nature, 388(6639), 279–284. https://doi.org/10.1038/40870

Evans, R., & Blackwell, K. (2015). Calcium: Amplitude, Duration, or Location? The Biological Bulletin, 228(1), 75. https://doi.org/10.1086/BBLV228N1P75

Eyal, G., Verhoog, M. B., Testa-Silva, G., Deitcher, Y., Benavides-Piccione, R., DeFelipe, J., de Kock, C. P. J., Mansvelder, H. D., & Segev, I. (2018). Human Cortical Pyramidal Neurons: From Spines to Spikes via Models. Frontiers in Cellular Neuroscience, 12(June), 1–24. https://doi.org/10.3389/fncel.2018.00181

Eyal, G., Verhoog, M. B., Testa-Silva, G., Deitcher, Y., Piccione, R. B., DeFelipe, J., de Kock, C. P. J., Mansvelder, H. D., & Segev, I. (2018). Human cortical pyramidal neurons: From spines to spikes via models. Frontiers in Cellular Neuroscience, 12, 181. https://doi.org/10.3389/FNCEL.2018.00181/BIBTEX

Fino, E., Paille, V., Cui, Y., Morera-Herreras, T., Deniau, J.-M., & Venance, L. (2010). Distinct coincidence detectors govern the corticostriatal spike timing-dependent plasticity. J Physiol, 588, 3045–3062. https://doi.org/10.1113/jphysiol.2010.188466

Fisher, R. E., Gray, R., & Johnston, D. (1990). Properties and Distribution of Single Voltage-Gated Calcium Channels in Adult Hippocampal Neurons. JOURNALOFNEUROPHYSIOLOGY, 64(1).

Frey, U., & Morris, R. G. M. (1997). Synaptic tagging and long-term potentiation. Nature, 385(6616), 533–536. https://doi.org/10.1038/385533A0

Gidon, A., & Segev, I. (2012). Principles Governing the Operation of Synaptic Inhibition in Dendrites. Neuron, 75(2), 330–341. https://doi.org/10.1016/j.neuron.2012.05.015

Golding, N. L., Kath, W. L., & Spruston, N. (2001). Dichotomy of Action-Potential Backpropagation in CA1 Pyramidal Neuron Dendrites. https://doi.org/10.1152/jn

Golding, N. L., Staff, N. P., & Spruston, N. (2002). Dendritic spikes as a mechanism for cooperative long-term potentiation. Nature, 418(6895), 326–331. https://doi.org/10.1038/nature00854

Graupner, M., & Brunel, N. (2012). Calcium-based plasticity model explains sensitivity of synaptic changes to spike pattern, rate, and dendritic location. Proceedings of the National Academy of Sciences of the United States of America, 109(10), 3991–3996. https://doi.org/10.1073/pnas.1109359109

Hardie, J., & Spruston, N. (2009). Synaptic Depolarization Is More Effective than Back-Propagating Action Potentials during Induction of Associative Long-Term Potentiation in Hippocampal Pyramidal Neurons. Journal of Neuroscience, 29(10), 3233–3241. https://doi.org/10.1523/JNEUROSCI.6000-08.2009

Hay, E., Hill, S., Schürmann, F., Markram, H., & Segev, I. (2011). Models of neocortical layer 5b pyramidal cells capturing a wide range of dendritic and perisomatic active properties. PLoS Computational Biology, 7(7). https://doi.org/10.1371/journal.pcbi.1002107

Hebb, D. (1949). The organization of behavior. A neuropsychological theory. https://pure.mpg.de/rest/items/item_2346268/component/file_2346267/content

Hines, M. L., Davison, A. P., & Muller, E. (2009). NEURON and Python. Frontiers in Neuroinformatics, 3(January), 1. https://doi.org/10.3389/neuro.11.001.2009

Humeau, Y., & Choquet, D. (2019). The next generation of approaches to investigate the link between synaptic plasticity and learning. Nature Neuroscience 2019 22:10, 22(10), 1536–1543. https://doi.org/10.1038/s41593-019-0480-6

Jadi, M. P., Behabadi, B. F., Poleg-Polsky, A., Schiller, J., & Mel, B. W. (2014). An augmented two-layer model captures nonlinear analog spatial integration effects in pyramidal neuron dendrites. Proceedings of the IEEE, 102(5), 782–798. https://doi.org/10.1109/JPROC.2014.2312671

Jadi, M., Polsky, A., Schiller, J., & Mel, B. W. (2012). Location-dependent effects of inhibition on local spiking in pyramidal neuron dendrites. PLoS Computational Biology, 8(6), e1002550. https://doi.org/10.1371/journal.pcbi.1002550

Jo, J., Heon, S., Kim, M. J., Son, G. H., Park, Y., Henley, J. M., Weiss, J. L., Sheng, M., Collingridge, G. L., & Cho, K. (2008). Metabotropic Glutamate Receptor-Mediated LTD Involves Two Interacting Ca2+ Sensors, NCS-1 and PICK1. Neuron, 60(6), 1095. https://doi.org/10.1016/J.NEURON.2008.10.050

Jones, I. S., & Kording, K. P. (2021). Might a Single Neuron Solve Interesting Machine Learning Problems Through Successive Computations on Its Dendritic Tree? Neural Computation, 33(6), 1554–1571. https://doi.org/10.1162/NECO_A_01390

Koch, C., Poggio, T., & Torre, V. (1983). Nonlinear interactions in a dendritic tree: Localization, timing, and role in information processing. Proceedings of the National Academy of Sciences of the United States of America, 80(9 I), 2799–2802. https://doi.org/10.1073/PNAS.80.9.2799

Koch, C., & Segev, I. (2000). The role of single neurons in information processing. Nature Neuroscience, 3(11s), 1171–1177. https://doi.org/10.1038/81444

Konur, S., Rabinowitz, D., Fenstermaker, V. L., & Yuste, R. (2003). Systematic regulation of spine sizes and densities in pyramidal neurons. Journal of Neurobiology, 56(2), 95–112. https://doi.org/10.1002/NEU.10229

Kumar, A., Barkai, E., & Schiller, J. (2021). Plasticity of olfactory bulb inputs mediated by dendritic NMDA-spikes in rodent piriform cortex. ELife, 10. https://doi.org/10.7554/ELIFE.70383

Landau, A. T., Park, P., Wong-Campos, J. D., Tian, H., Cohen, A. E., & Sabatini, B. L. (2022). Dendritic branch structure compartmentalizes voltage-dependent calcium influx in cortical layer 2/3 pyramidal cells. ELife, 11. https://doi.org/10.7554/ELIFE.76993

Larkum, M. E. (2022). Are Dendrites Conceptually Useful? Neuroscience, 489, 4–14. https://doi.org/10.1016/J.NEUROSCIENCE.2022.03.008

Lisman, J. (1989). A mechanism for the Hebb and the anti-Hebb processes underlying learning and memory. Proceedings of the National Academy of Sciences, 86(23), 9574–9578. https://doi.org/10.1073/pnas.86.23.9574

Lisman, J. E. (2001). Three Ca2+ levels affect plasticity differently: the LTP zone, the LTD zone and no man’s land. The Journal of Physiology, 532(2), 285–285. https://doi.org/10.1111/J.1469-7793.2001.0285F.X

Lisman, J., & Spruston, N. (2005). Postsynaptic depolarization requirements for LTP and LTD: a critique of spike timing-dependent plasticity. Nature Neuroscience, 8(7), 839–841. https://doi.org/10.1038/nn0705-839

Liu, H., Lan, Y., Bing, Y. H., Chu, C. P., & Qiu, D. L. (2016). N-methyl-D-Aspartate Receptors Contribute to Complex Spike Signaling in Cerebellar Purkinje Cells: An In vivo Study in Mice. Frontiers in Cellular Neuroscience, 10(Jun). https://doi.org/10.3389/FNCEL.2016.00172

Lynch, G. S., Dunwiddie, T., & Gribkoff, V. (1977). Heterosynaptic depression: a postsynaptic correlate of long-term potentiation. Nature, 266(5604), 737–739. https://doi.org/10.1038/266737a0

Malenka, R. C., Kauer, J. A., Perkel, D. J., Mauk, M. D., Kelly, P. T., Nicoll, R. A., & Waxham, M. N. (1989). An essential role for postsynaptic calmodulin and protein kinase activity in long-term potentiation. 340(6234), 554–557. https://www.nature.com/articles/340554a0

Malinow, R., Schulman, H., & Tsien, R. W. (1989). Inhibition of postsynaptic PKC or CaMKII blocks induction but not expression of LTP. Science (New York, N.Y.), 245(4920), 862–866. https://doi.org/10.1126/SCIENCE.2549638

Mel, B. W. (1991). The clusteron: Toward a simple abstraction for a complex neuron. Nips, 35–42.

Moldwin, T., Kalmenson, M., & Segev, I. (2021). The gradient clusteron: A model neuron that learns to solve classification tasks via dendritic nonlinearities, structural plasticity, and gradient descent. PLOS Computational Biology, 17(5), e1009015. https://doi.org/10.1371/journal.pcbi.1009015

Moldwin, T., & Segev, I. (2020). Perceptron Learning and Classification in a Modeled Cortical Pyramidal Cell. Frontiers in Computational Neuroscience, 14(April), 1–13. https://doi.org/10.3389/fncom.2020.00033

Mulkey, R. M., Endo, S., Shenolikar, S., & Malenka, R. C. (1994). Involvement of a calcineurin/ inhibitor-1 phosphatase cascade in hippocampal long-term depression. Nature 1994 369:6480, 369(6480), 486–488. https://doi.org/10.1038/369486a0

Mulkey, R. M., Herron, C. E., & Malenka, R. C. (1993). An essential role for protein phosphatases in hippocampal long-term depression. Science (New York, N.Y.), 261(5124), 1051–1055. https://doi.org/10.1126/SCIENCE.8394601

Mulkey, R. M., & Malenka, R. C. (1992). Mechanisms underlying induction of homosynaptic long-term depression in area CA1 of the hippocampus. Neuron, 9(5), 967–975. https://doi.org/10.1016/0896-6273(92)90248-C

Nabavi, S., Fox, R., Proulx, C. D., Lin, J. Y., Tsien, R. Y., & Malinow, R. (2014). Engineering a memory with LTD and LTP. Nature, 511(7509), 348–352. https://doi.org/10.1038/nature13294

Neveu, D., & Zucker, R. S. RS. (1996). Postsynaptic Levels of [Ca2+]i Needed to Trigger LTD and LTP. Neuron, 16(3), 619–629. https://doi.org/10.1016/S0896-6273(00)80081-1

Nishiyama, M., Hong, K., Mikoshiba, K., Poo, M. M., & Kato, K. (2000). Calcium stores regulate the polarity and input specificity of synaptic modification. Nature, 408(6812), 584–588. https://doi.org/10.1038/35046067

O’Connor, D. H., Wittenberg, G. M., & Wang, S. S. H. (2005a). Graded bidirectional synaptic plasticity is composed of switch-like unitary events. Proceedings of the National Academy of Sciences of the United States of America, 102(27), 9679–9684. https://doi.org/10.1073/PNAS.0502332102

O’Connor, D. H., Wittenberg, G. M., & Wang, S. S.-H. (2005b). Dissection of Bidirectional Synaptic Plasticity Into Saturable Unidirectional Processes. Journal of Neurophysiology, 94(2), 1565–1573. https://doi.org/10.1152/jn.00047.2005

O’Hare, J. K., Gonzalez, K. C., Herrlinger, S. A., Hirabayashi, Y., Hewitt, V. L., Blockus, H., Szoboszlay, M., Rolotti, S. v., Geiller, T. C., Negrean, A., Chelur, V., Polleux, F., & Losonczy, A. (2022). Compartment-specific tuning of dendritic feature selectivity by intracellular Ca2+ release. Science, 375(6586). https://doi.org/10.1126/science.abm1670

Palmer, L. M., Shai, A. S., Reeve, J. E., Anderson, H. L., Paulsen, O., & Larkum, M. E. (2014). NMDA spikes enhance action potential generation during sensory input. Nature Neuroscience, 17(3), 383–390. https://doi.org/10.1038/NN.3646

Piochon, C., Irinopoulou, T., Brusciano, D., Bailly, Y., Mariani, J., & Levenes, C. (2007). NMDA Receptor Contribution to the Climbing Fiber Response in the Adult Mouse Purkinje Cell. Journal of Neuroscience, 27(40), 10797–10809. https://doi.org/10.1523/JNEUROSCI.2422-07.2007

Piochon, C., Levenes, C., Ohtsuki, G., & Hansel, C. (2010). Purkinje cell NMDA receptors assume a key role in synaptic gain control in the mature cerebellum. The Journal of Neuroscience: The Official Journal of the Society for Neuroscience, 30(45), 15330–15335. https://doi.org/10.1523/JNEUROSCI.4344-10.2010

Piochon, C., Titley, H. K., Simmons, D. H., Grasselli, G., Elgersma, Y., & Hansel, C. (2016). Calcium threshold shift enables frequency-independent control of plasticity by an instructive signal. 46, 13221–13226. https://www.pnas.org/content/113/46/13221

Poirazi, P., Brannon, T., & Mel, B. W. (2003). Pyramidal Neuron as Two-Layer Neural Network. Neuron, 37(6), 989–999. https://doi.org/10.1016/S0896-6273(03)00149-1

Poirazi, P., & Mel, B. W. (2001). Impact of active dendrites and structural plasticity on the memory capacity of neural tissue. Neuron, 29(3), 779–796. https://doi.org/10.1016/S0896-6273(01)00252-5

Poleg-Polsky, A. (2015). Effects of neural morphology and input distribution on synaptic processing by global and focal NMDA-spikes. PLoS ONE, 10(10). https://doi.org/10.1371/JOURNAL.PONE.0140254

Polsky, A., Mel, B., & Schiller, J. (2009). Encoding and decoding bursts by NMDA spikes in basal dendrites of layer 5 pyramidal neurons. Journal of Neuroscience, 29(38), 11891–11903. https://doi.org/10.1523/JNEUROSCI.5250-08.2009

Polsky, A., Mel, B. W., & Schiller, J. (2004). Computational subunits in thin dendrites of pyramidal cells. Nature Neuroscience, 7(6), 621–627. https://doi.org/10.1038/nn1253

Rall, W. (1967). Distinguishing theoretical synaptic potentials computed for different soma-dendritic distributions of synaptic input. Journal of Neurophysiology, 30(5), 1138–1168. https://doi.org/10.1152/jn.1967.30.5.1138

Rall, W., & Rinzel, J. (1973). Branch input resistance and steady attenuation for input to one branch of a dendritic neuron model. Biophysical Journal, 13(7), 648–688. https://doi.org/10.1016/S0006-3495(73)86014-X

Redondo, R. L., Okuno, H., Spooner, P. A., Frenguelli, B. G., Bito, H., & Morris, R. G. M. (2010). Synaptic Tagging and Capture: Differential Role of Distinct Calcium/Calmodulin Kinases in Protein Synthesis-Dependent Long-Term Potentiation. Journal of Neuroscience, 30(14), 4981–4989. https://doi.org/10.1523/JNEUROSCI.3140-09.2010

Rodrigues, Y. E., Tigaret, C. M., Marie, H., O’Donnell, C., & Veltz, R. (2021). A stochastic model of hippocampal synaptic plasticity with geometrical readout of enzyme dynamics. BioRxiv, 2021.03.30.437703. https://doi.org/10.1101/2021.03.30.437703

Rose, C. R., & Konnerth, A. (2001). Stores Not Just for Storage: Intracellular Calcium Release and Synaptic Plasticity. Neuron, 31(4), 519–522. https://doi.org/10.1016/S0896-6273(01)00402-0

Rose, G. M., & Dunwiddie, T. v. (1986). Induction of hippocampal long-term potentiation using physiologically patterned stimulation. Neuroscience Letters, 69(3), 244–248. https://doi.org/10.1016/0304-3940(86)90487-8

Royer, S., & Paré, D. (2003). Conservation of total syanptic wight through balanced synaptic depression and potentiation. Nature, 422(April), 518–522. https://doi.org/10.1038/nature01532.1.

Rumelhart, D. E., Hinton, G. E., & Williams, R. J. (1986). Learning representations by back-propagating errors. Nature, 323(6088), 533–536. https://doi.org/10.1038/323533a0

Sabatini, B. L., Oertner, T. G., & Svoboda, K. (2002). The Life Cycle of Ca2+ Ions in Dendritic Spines. Neuron, 33(3), 439–452. https://doi.org/10.1016/S0896-6273(02)00573-1

Segev, I., & Rall, W. (1988). Computational study of an excitable dendritic spine. Journal of Neurophysiology, 60(2), 499–523. https://doi.org/10.1152/jn.1988.60.2.499

Segev, I., & Rall, W. (1998). Excitable dendrites and spines: earlier theoretical insights elucidate recent direct observations. Trends in Neurosciences, 21(11), 453–460. https://doi.org/10.1016/S0166-2236(98)01327-7

Shindou, T., Ochi-Shindou, M., & Wickens, J. R. (2011). A Ca2+ Threshold for Induction of Spike-Timing-Dependent Depression in the Mouse Striatum. Journal of Neuroscience, 31(36), 13015–13022. https://doi.org/10.1523/JNEUROSCI.3206-11.2011

Shouval, H. Z., Bear, M. F., & Cooper, L. N. (2002). A unified model of NMDA receptor-dependent bidirectional synaptic plasticity. Proceedings of the National Academy of Sciences, 99(16), 10831–10836. https://doi.org/10.1073/PNAS.152343099

Shouval, H. Z., Wang, S. S. H.-H., & Wittenberg, G. M. (2010). Spike timing dependent plasticity: A consequence of more fundamental learning rules. Frontiers in Computational Neuroscience, 4(1), 19. https://doi.org/10.3389/fncom.2010.00019

Snutch, T. P., Peloquin, J., Mathews, E., & McRory, J. E. (2013). Molecular Properties of Voltage-Gated Calcium Channels. https://www.ncbi.nlm.nih.gov/books/NBK6181/

Sun, C., Nold, A., Fusco, C. M., Rangaraju, V., Tchumatchenko, T., Heilemann, M., & Schuman, E. M. (2021). The prevalence and specificity of local protein synthesis during neuronal synaptic plasticity. Science Advances, 7(38), 790–807. https://doi.org/10.1126/SCIADV.ABJ0790/SUPPL_FILE/SCIADV.ABJ0790_SM.PDF

Tigaret, C. M., Olivo, V., Sadowski, J. H. L. P., Ashby, M. C., & Mellor, J. R. (2016). Coordinated activation of distinct Ca2+ sources and metabotropic glutamate receptors encodes Hebbian synaptic plasticity. Nature Communications, 7. https://doi.org/10.1038/ncomms10289

Tong, R., Chater, T. E., Emptage, N. J., & Goda, Y. (2021). Heterosynaptic cross-talk of pre-and postsynaptic strengths along segments of dendrites. Cell Reports, 34(4), 108693. https://doi.org/10.1016/J.CELREP.2021.108693

Tsien, R. W., Lipscombe, D., Madison, D. v., Bley, K. R., & Fox, A. P. (1988). Multiple types of neuronal calcium channels and their selective modulation. Trends in Neurosciences, 11(10), 431–438. https://doi.org/10.1016/0166-2236(88)90194-4

White, G., Levy, W. B., & Steward, O. (1988). Evidence that associative interactions between synapses during the induction of long-term potentiation occur within local dendritic domains. Proceedings of the National Academy of Sciences, 85(7), 2368–2372. https://doi.org/10.1073/PNAS.85.7.2368

Whitlock, J., Heynen, A., Shuler, M., & Bear, M. (2006). Learning induces long-term potentiation in the hippocampus. Science, 313, 1093–1097.

Yang, S., Tang, Y., Zucker, R. S., Yang, S.-N., Tang, Y.-G., & Zucker Selective, R. S. (1999). Selective Induction of LTP and LTD by Postsynaptic [Ca 2] i Elevation.

Yuste, R., & Denk, W. (1995). Dendritic spines as basic functional units of neuronal integration. Nature, 375(6533), 682–684. https://doi.org/10.1038/375682a0

